# Global trends in biodiversity and ecosystem services from 1900 to 2050

**DOI:** 10.1101/2020.04.14.031716

**Authors:** Henrique M. Pereira, Isabel M.D. Rosa, Inês S. Martins, HyeJin Kim, Paul Leadley, Alexander Popp, Detlef P. van Vuuren, George Hurtt, Peter Anthoni, Almut Arneth, Daniele Baisero, Rebecca Chaplin-Kramer, Louise Chini, Fulvio Di Fulvio, Moreno Di Marco, Simon Ferrier, Shinichiro Fujimori, Carlos A. Guerra, Michael Harfoot, Thomas D. Harwood, Tomoko Hasegawa, Vanessa Haverd, Petr Havlík, Stefanie Hellweg, Jelle P. Hilbers, Samantha L. L. Hill, Akiko Hirata, Andrew J. Hoskins, Florian Humpenöder, Jan H. Janse, Walter Jetz, Justin A. Johnson, Andreas Krause, David Leclère, Tetsuya Matsui, Johan R. Meijer, Cory Merow, Michael Obsersteiner, Haruka Ohashi, Benjamin Poulter, Andy Purvis, Benjamin Quesada, Carlo Rondinini, Aafke M. Schipper, Josef Settele, Richard Sharp, Elke Stehfest, Bernardo B. N. Strassburg, Kiyoshi Takahashi, Matthew V. Talluto, Wilfried Thuiller, Nicolas Titeux, Piero Visconti, Christopher Ware, Florian Wolf, Rob Alkemade

**Affiliations:** German Centre for Integrative Biodiversity Research (iDiv) Halle-Jena-Leipzig, Deutscher Platz 5e, 04103 Leipzig, Germany; Institute of Biology, Martin Luther University Halle Wittenberg, Am Kirchtor 1, 06108 Halle (Saale), Germany; CIBIO/InBIO, Centro de Investigação em Biodiversidade e Recursos Genéticos, Universidade do Porto, Campus Agrário de Vairão, R. Padre Armando Quintas, 4485-661 Vairão, Portugal; School of Natural Sciences, Bangor University, Bangor, Gwynedd, LL57 2DG; Ecologie Systématique Evolution, Univ. Paris-Sud, CNRS, AgroParisTech, Université Paris-Saclay, 91400, Orsay, France; Potsdam Institute for Climate Impact Research (PIK), Member of the Leibniz Association, Potsdam, Germany; PBL Netherlands Environmental Assessment Agency, the Hague, the Netherlands; Copernicus Institute for Sustainable Development, Utrecht University, Utrecht, the Netherlands; Department of Geographical Sciences, University of Maryland, College Park, MD 20742, USA; Karlsruhe Institute of Technology, Dept. Meteorology and Climate/Atmospheric Environmental Research, Kreuzeckbahnstr. 19, 82467 Garmisch-Partenkirchen, Germany; Global Mammal Assessment program, Department of Biology and Biotechnologies, Sapienza Università di Roma, Viale dell’Università 32, I-00185, Rome, Italy; The Natural Capital Project, Stanford University, 371 Serra Mall, Stanford, CA 94305, USA; International Institute for Applied Systems Analysis, Schlossplatz 1, Laxenburg 2361, Austria; CSIRO Land and Water, GPO Box 2583, Brisbane QLD 4001, Australia; CSIRO Land and Water, GPO Box 1700, Canberra ACT 2601, Australia; Kyoto University, Department of Environmental Engineering, 361, C1-3, Kyoto University Katsura Campus, Nishikyo-ku, Kyoto-city, 615-8540 Japan; Center for Climate Change Adaptation, National Institute for Environmental Studies (NIES), 16-2 Onogawa, Tsukuba, Ibaraki 305-8506, Japan; UN Environment, World Conservation Monitoring Centre, 219 Huntingdon Road, Cambridge, CB3 0DL, UK; College of Science and Engineering, Ritsumeikan University, 1-1-1, Nojihigashi, Kusatsu, Shiga, 525-8577, Japan; CSIRO Oceans and Atmosphere, Canberra, 2601, Australia; Institute of Environmental Engineering, ETH Zurich, 8093 Zurich, Switzerland; Department of Life Sciences, Natural History Museum, London SW7 5BD, UK; Forestry and Forest Products Research Institute, Forest Research and Management Organization, 1 Matsunosato, Tsukuba, Ibaraki, 305-8687, Japan; CSIRO Health and Biosecurity, James Cook University, Townsville, QLD 4810, Australia; Netherlands Inst. of Ecology NIOO-KNAW, Wageningen, the Netherlands; Ecology and Evolutionary Biology, Yale University, 165 Prospect St, New Haven, CT 06511, USA; Natural Capital Project, Institute on the Environment, University of Minnesota, 1954 Buford Ave, Saint Paul, MN 55108, USA; Technical University of Munich, TUM School of Life Sciences Weihenstephan, 85354 Freising, Germany; Ecology and Evolutionary Biology, University of Connecticut, Unit 3043, Storrs CT 06269; Environmental Change Institute, South Parks Road, Oxford, OX1 3QY, United Kingdom; NASA Goddard Space Flight Center, Biospheric Sciences Laboratory, Greenbelt, MD 20771, USA; Department of Life Sciences, Imperial College London, Silwood Park, Ascot SL5 7PY, UK; Universidad del Rosario, Faculty of Natural Sciences and Mathematics, Kr 26 No 63B-48, Bogotá D.C, Colombia; Radboud University, Institute for Water and Wetland Research, PO Box 9010, 6500 GL Nijmegen, Netherlands; Helmholtz Centre for Environmental Research – UFZ, Department of Community Ecology, Theodor-Lieser-Strasse 4, 06210 Halle, Germany; Institute of Biological Sciences, University of the Philippines, Los Baños, College, 4031, Laguna, Philippines; Rio Conservation and Sustainability Science Centre, Department of Geography and the 10 Environment, Pontifícia Universidade Católica, 22453900, Rio de Janeiro, Brazil; International Institute for Sustainability, Estrada Dona Castorina 124, 22460-320, Rio de Janeiro, Brazil; Postgraduate Programme in Ecology, Universidade Federal do Rio de Janeiro, 21941-590, Rio de Janeiro, Brazil; Leibniz-Institute of Freshwater Ecology and Inland Fisheries (IGB), Müggelseedamm 310, 12587 Berlin, Germany; Univ. Grenoble Alpes, CNRS, Univ. Savoie Mont Blanc, LECA, Laboratoire d’Écologie Alpine, F-38000 Grenoble, France; Université Catholique de Louvain, Earth and Life Institute, Biodiversity Research Centre, 1348 Louvain-la-Neuve, Belgium; Institute of Zoology, Zoological Society of London, Regent’s Park, London, NW1 4RY; Centre for Biodiversity and Environment Research, University College London, Gower Street, London, C1E6BT, UK; Environmental System Analysis Group, Wageningen University, Wageningen, Netherlands

## Abstract

Despite the scientific consensus on the extinction crisis and its anthropogenic origin, the quantification of historical trends and of future scenarios of biodiversity and ecosystem services has been limited, due to the lack of inter-model comparisons and harmonized scenarios. Here, we present a multi-model analysis to assess the impacts of land-use and climate change from 1900 to 2050. During the 20th century provisioning services increased, but biodiversity and regulating services decreased. Similar trade-offs are projected for the coming decades, but they may be attenuated in a sustainability scenario. Future biodiversity loss from land-use change is projected to keep up with historical rates or reduce slightly, whereas losses due to climate change are projected to increase greatly. Renewed efforts are needed by governments to meet the 2050 vision of the Convention on Biological Diversity.

**One Sentence Summary:** Development pathways exist that allow for a reduction of the rates of biodiversity loss from land-use change and improvement in regulating services but climate change poses an increasing challenge.

## Main Text

During the last century humans have caused biodiversity loss at rates higher than ever before, with extinction rates for vertebrates of 0.5% to 1% per century, 50 to100 times higher than the mean extinction rates in the Cenozoic fossil record (*1*–*4*). Although the proximate causes of this loss are multiple, ultimately a growing human population and economy have led to an increasing demand for land and natural resources causing habitat conversion and loss (*5*).

Associated increases in the flow of provisioning ecosystem services such as the production of crops and livestock also lead to the widespread degradation of ecosystem’s capacity to provide regulating services such as pollination and water quality, raising concerns about the long-term sustainability of recent development trends (*6*). Addressing the biodiversity crisis is increasingly at the center of international policy-making, under multilateral agreements such as the Convention on Biological Diversity. Restoring biodiversity and ecosystem services can actually provide part of the solution to many of the UN Sustainable Development Challenges (*7, 8*). Therefore, it is key to assess implications of future socio-economic developments for biodiversity and ecosystem services and identify policies that can shift developments towards more sustainable pathways.

Scenario studies examine alternative future socio-economic development pathways and their impacts on direct drivers such as land-use and climate, often using integrated assessment models (*9*). The scenarios consequences for biodiversity and ecosystem services can be assessed using biodiversity and ecosystem function and services models (*10, 11*). Several studies have explored the future trends of biodiversity and ecosystem services, finding an acceleration of extinction rates 100 to 10 000 times higher than the fossil record, and the continuation of trends of increasing provisioning services with the degradation of some regulation services, although with strong regional variations (*10, 12*–*15*). While enlightening on the potential trajectories of biodiversity under global changes, these studies are hardly comparable. Existing scenario studies often use a single model, analyze a single facet of biodiversity, lack integration between biodiversity and ecosystem services impacts, or when comparing multiple models use different projections for future land-use and climate. Therefore, the source of uncertainties in these scenarios is difficult to ascertain (*16*) and an integrated analysis of biodiversity and ecosystem services scenarios has remained elusive.

Here, we present the first multi-model ensemble projections of biodiversity and ecosystem services using a set of harmonized land-use and climate change reconstructions from 1900 to 2015 and three future scenarios from 2015 to 2050. This work was carried out under the auspices of the Expert Group on Scenarios and Models of the Intergovernmental Platform on Biodiversity and Ecosystem Services (IPBES) (*17*). We quantified a set of common ecological metrics from the grid cell scale (*α*-metrics), to the regional (i.e., IPBES subregions) and global scale (*γ*-metrics) to answer two main questions: (1) What are the global impacts of land-use and climate change on multiple facets of biodiversity and ecosystem services (i.e., Nature’s Contributions to People, NCP) over the coming decades, compared to their impacts during the 20th century? (2) How much of the variation in projected impacts can be attributed to differences of development pathways in scenarios and to differences between models (i.e. structural uncertainty)?

We explored a range of plausible futures using the scenario framework of the Shared Socio-Economic Pathways (SSP) and Representative Concentration Pathways (RCP) (*18*). We chose three specific SSP-RCP combinations representing different storylines of population growth, socio-economic development and the level of greenhouse gas emissions (climate policy). These combinations represent contrasting projections of future land-use and climate change (Table S1, Figures S1 and S2): SSP1xRCP2.6 (“global sustainability” with low climate change and low land-use change), SSP3xRCP6.0 (“regional rivalry” with intermediate climate change and high land-use change) and SSP5xRCP8.5 (“fossil-fueled development” with high climate change and intermediate land-use change). For the biodiversity analysis, we consider both the impacts of land-use change alone (maintaining climate constant at historical levels) and of land-use change and climate change combined.

We brought together eight models of biodiversity and five models of ecosystem function and services (Table S2). Depending on the model, up to three biodiversity metrics were calculated (SM): species richness (S), mean species habitat score 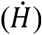, and species-abundance based biodiversity intactness (I). For ecosystem functions and services, we classified model outputs into nine classes of Nature’s Contributions to People (*19*) (Table S1). We calculated the metrics at the grid cell level (*α*), at the regional level, and at the global level (*γ*).

The steep reduction in global species richness that occurred during the 20^th^ century (−0.78% ± 0.30% per century, mean±SE across models) is expected to continue at a slower (global sustainability scenario) or at a similar pace (regional rivalry and fossil-fueled development scenarios) in the next decades when land-use change alone is considered (Figure 1a). However, a much steeper decline is expected when the combined effects of land-use change and climate change are considered (Figure 1b, 1c). The scenario where we are able to stabilize greenhouse gas emissions concentrations and limit climate change to 2°C (global sustainability scenario; Figure S2) has already 40% lower global extinction rates by 2050 than the scenario with no climate mitigation policy (fossil-fueled development), with bigger differences looming for the second half of this century as the contrast between these scenarios continues to increase (*20*). Other biodiversity metrics exhibit similar trends with some interesting differences. Reduction in local species richness are of similar magnitude to global species richness changes, while biodiversity metrics based on global habitat extent across species or abundance-based intactness are up to an order of magnitude more sensitive to land-use change (Figure 1b). The uncertainties due to inter-model variation are large, particularly for the climate change impacts which are based on a smaller subset of models, but the trends are still clear (Figure 1b, 1c).

**Fig. 1.**
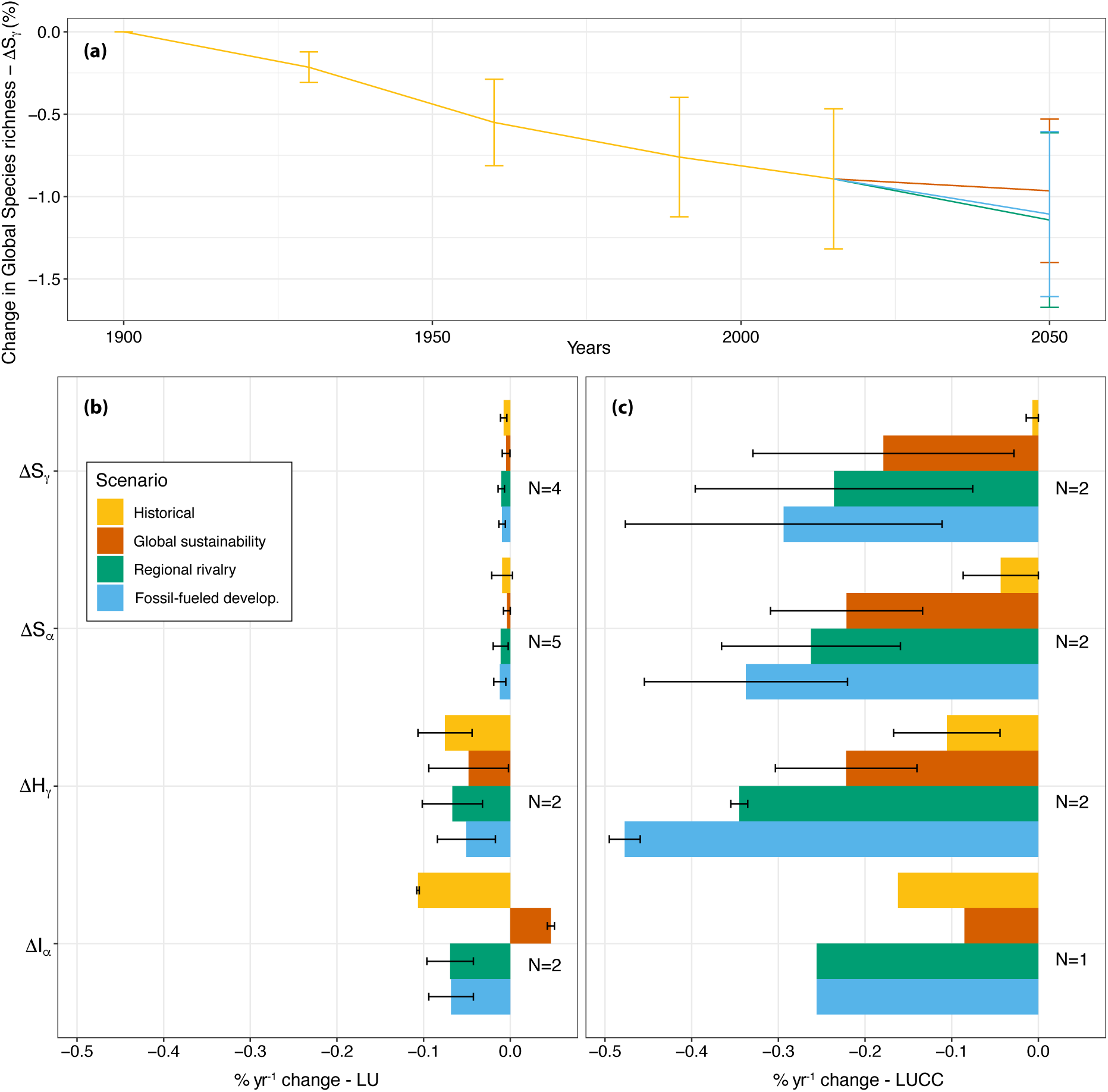
Historical trends in biodiversity since 1900 and future projections for each scenario to 2050. **(a)** Proportional global species richness change (ΔS_*γ*_) relative to 1900 from land-use change only. Change in different dimensions of biodiversity for the historical period (1900-2015) and for each future scenario (2015-2050): **(c)** from land-use alone; **(d)** from land-use change and climate change combined. Metrics correspond to proportional changes in: global species richness (Δ*S*_*γ*_), local species richness averaged across space 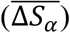, mean species global habitat extent 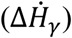, and local intactness averaged across space 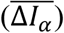. All values given as means across models with error bars representing standard errors. *N* is number of models used for metric.

Global averages mask some even larger species reductions at the level of individual grid-cells (Figure 2). During the 20^th^ century, net reductions in local species richness occurred across much of the world, with pronounced losses in Central America, the Andes, the Southeast of Brazil, West Africa, East Africa, South-East Asia, Eastern Australia and South-West Australia, Central North America, Madagascar, New Zealand and the Caribbean (Figure 2a). In the future, some of these regions, particularly in the tropics, are projected to see further biodiversity losses from land-use change (Figure 2b-d), while some regions start seeing losses for the first time, particularly in the Northern boreal regions as forestry activities increase, and regions in central Africa because of conversion to pasture (Figure S1e). In contrast, some areas in Western Europe and Northeast America have seen modest net gains in local species richness during the last century, as a result of farmland abandonment and decrease of forestry (Figure S1c) This pattern is expected to expand in the future to other temperate areas (Figure 2b-c). However, those regions already incurred extinctions before 1900 and these limited increases are not enough to noticeably improve biodiversity intactness (Figure S3).

**Fig. 2:**
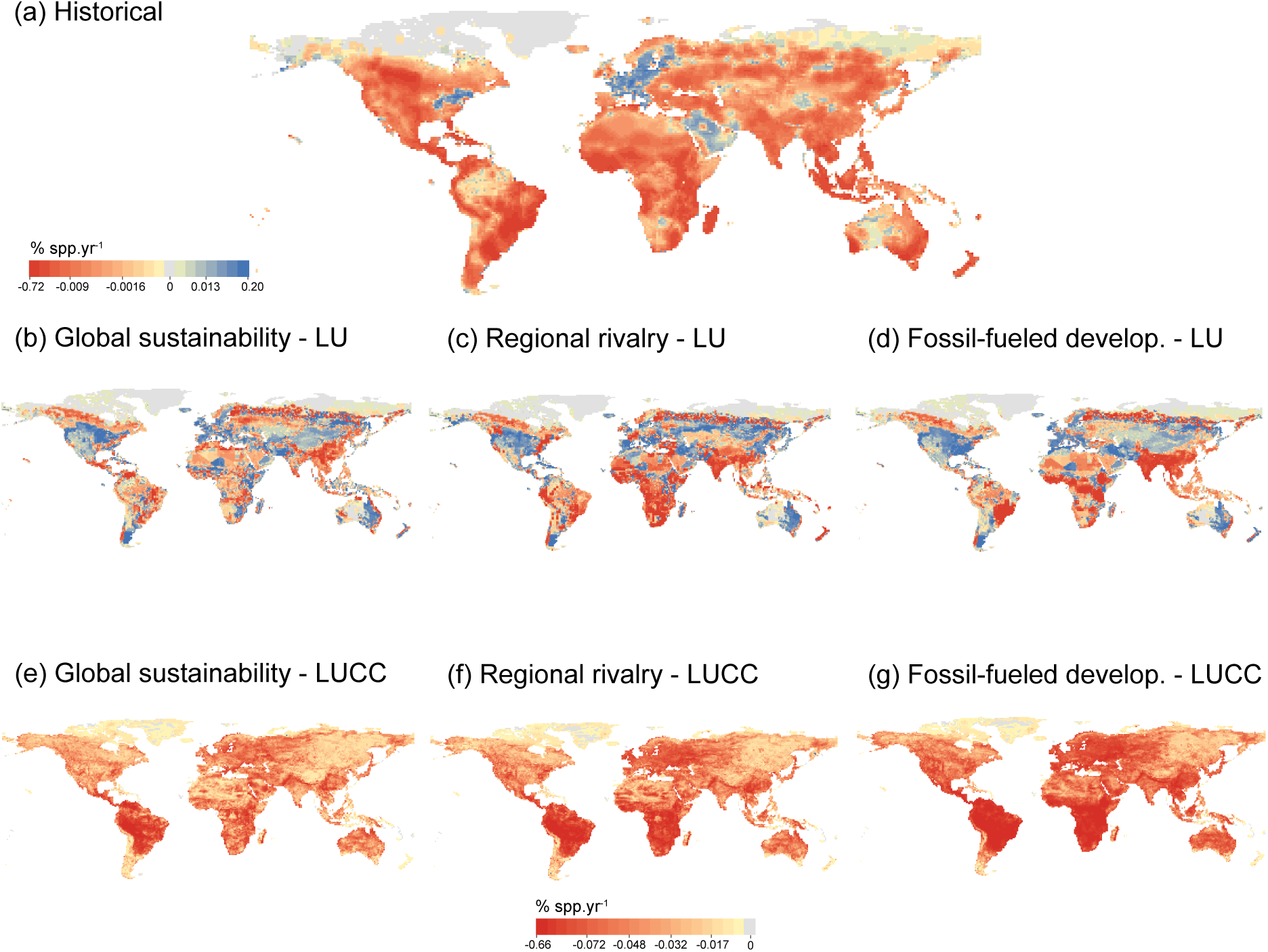
Spatial distribution of absolute changes in local species richness per year (*ΔSS*_*α*_). (a) Historical changes from 1900 to 2015 (number of models, N=5). Future projected changes 2015 to 2050 caused by land-use change alone in each scenario (**b-d**; N=5) and by land-use change and climate change combined (**e-f**, N=2). All values are based on inter-model means and normalized relative to the maximum local species richness in each model (e.g. a value of −50% corresponds to a reduction in species richness equal to half of the maximum species richness across cells). Color scale is based on quantile intervals and differs for **(a-d)** and **(e-g)**.

The three future scenarios exhibit important regional contrasts of biodiversity change. In the global sustainability scenario further land-use-induced losses are moderate and largely restricted to areas that have already been degraded in the last century (Figure 2b). In the regional rivalry scenario, a more regionalized socio-economic development leads to multiple fronts of biodiversity loss across the world including developed and developing regions (Figure 2c), while in the fossil-fueled development scenario a more globalized world sees biodiversity loss concentrated in Southeast South America, Central Africa, East Africa and South Asia (Figure 2d). When climate is also considered, the losses are further exacerbated: losses occur in much of the world, and especially concentrated in the highly biodiverse areas in the Neotropics and Afrotropics (Figure 2e-g). Spatial patterns are broadly consistent across models, although some disagreement exists, particularly regarding areas where local species richness may increase (Figure S4). When relative changes in species richness are compared with absolute changes (Figure S5), it is apparent that the latter are larger in tropical regions and continents (except Australia), as temperate areas and islands often have lower species richness.

During the 20^th^ century, increases material ecosystem services at the global scale, such as food and timber provisioning, were obtained at the cost of regulating services, such as pollination and nutrient retention (Figure 3). The same overall trends and trade-offs are projected for the next few decades, although much less pronounced in the global sustainability scenario, where limited population growth combined with healthy diets and reduction of food waste, leads to the smallest increases in food, feed and timber demand. This, associated with increases in agricultural productivity and other environmental policies, allows for improvements in some regulating ecosystem services and only moderate declines in others. The global sustainability scenario also has the largest increase in bioenergy production as a component of climate mitigation policies, which leads to land-use change (Figure S1a) and impacts on biodiversity (Figure 2b).

**Fig. 3:**
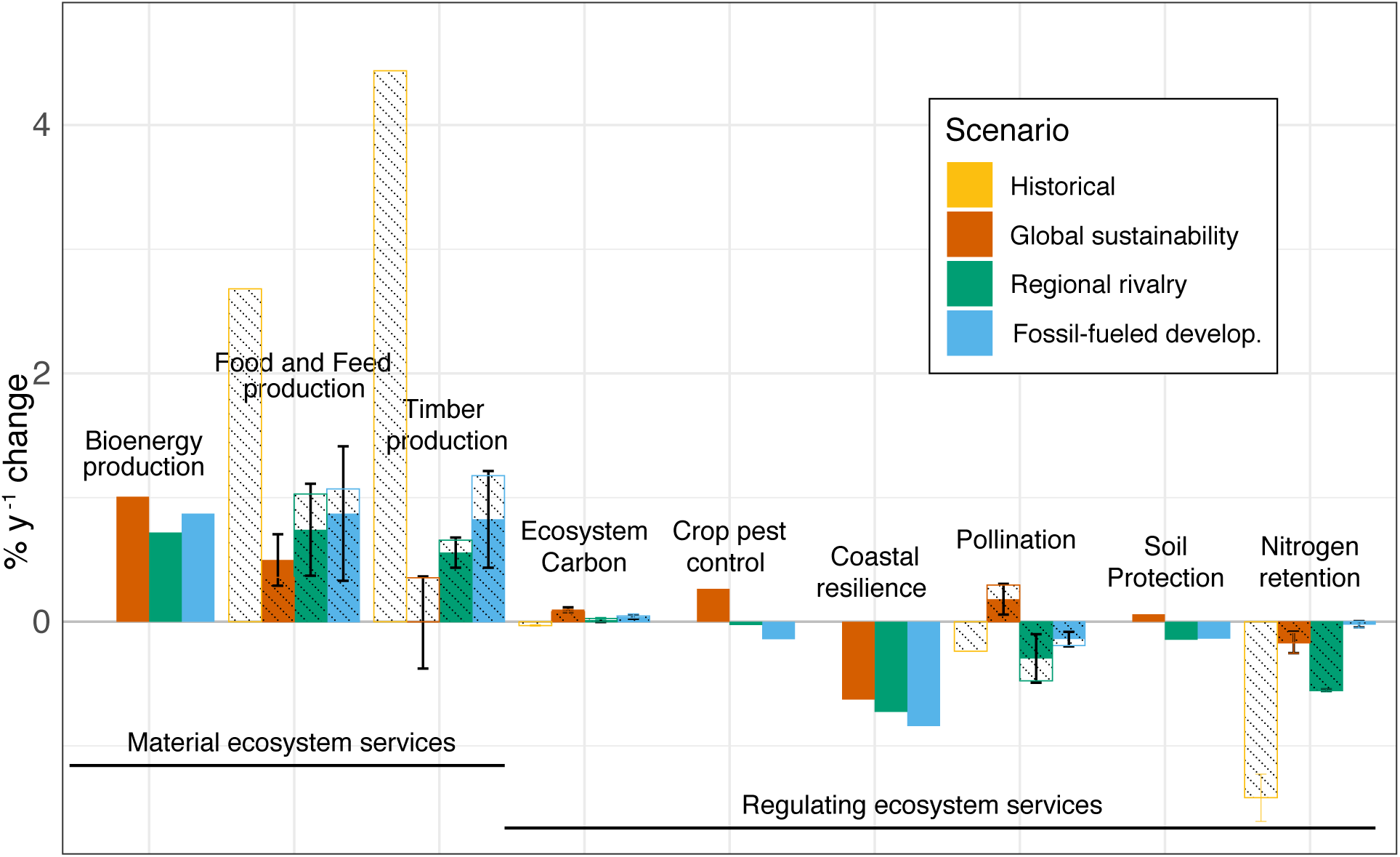
Historical (1900-2015) rate of changes in material and regulating ecosystem services at the global level and future projections for each scenario (2015-2050). For services assessed with more than one model, reported values are inter-model means and error bars represent standard errors. Dashed bars correspond to the subset of models that project historical changes.

In the two other scenarios, larger rates of increase in food and feed, and timber demand are projected (c. 1% yr^-1^), although smaller than during the last century (c. 3-4% yr^-1^) due to decelerating population growth, while decreases are projected for crop pest control, coastal resilience, pollination, soil protection, and nitrogen retention (Figure 3). In contrast with the biodiversity projections, the scenario with highest climate change (fossil fueled development) does not generally have more negative consequences for regulating services than the scenario with intermediate climate change (regional rivalry). The exception is that increasing climate change is likely to play a major role in increasing vulnerability of coastal populations.

Surprisingly, little change in total ecosystem carbon is anticipated between scenarios, probably due to CO_2_ fertilization effects in higher climate change scenarios (regional rivalry and fossil fueled development; Figure S6) compensating for the decreases in total forest area (Figure S1a). There is some inter-model variation in the projections of individual ecosystem services. Models for some ecosystem services exhibit strong spatial agreement, such as for ecosystem carbon (Figure S7), while for other ecosystem services, models still exhibit some regions of disagreement, such as for food and feed production (Figure S8). Still, in most cases regional or global variation between scenarios is greater than variation between models (Figure 3 and 4).

**Fig. 4.**
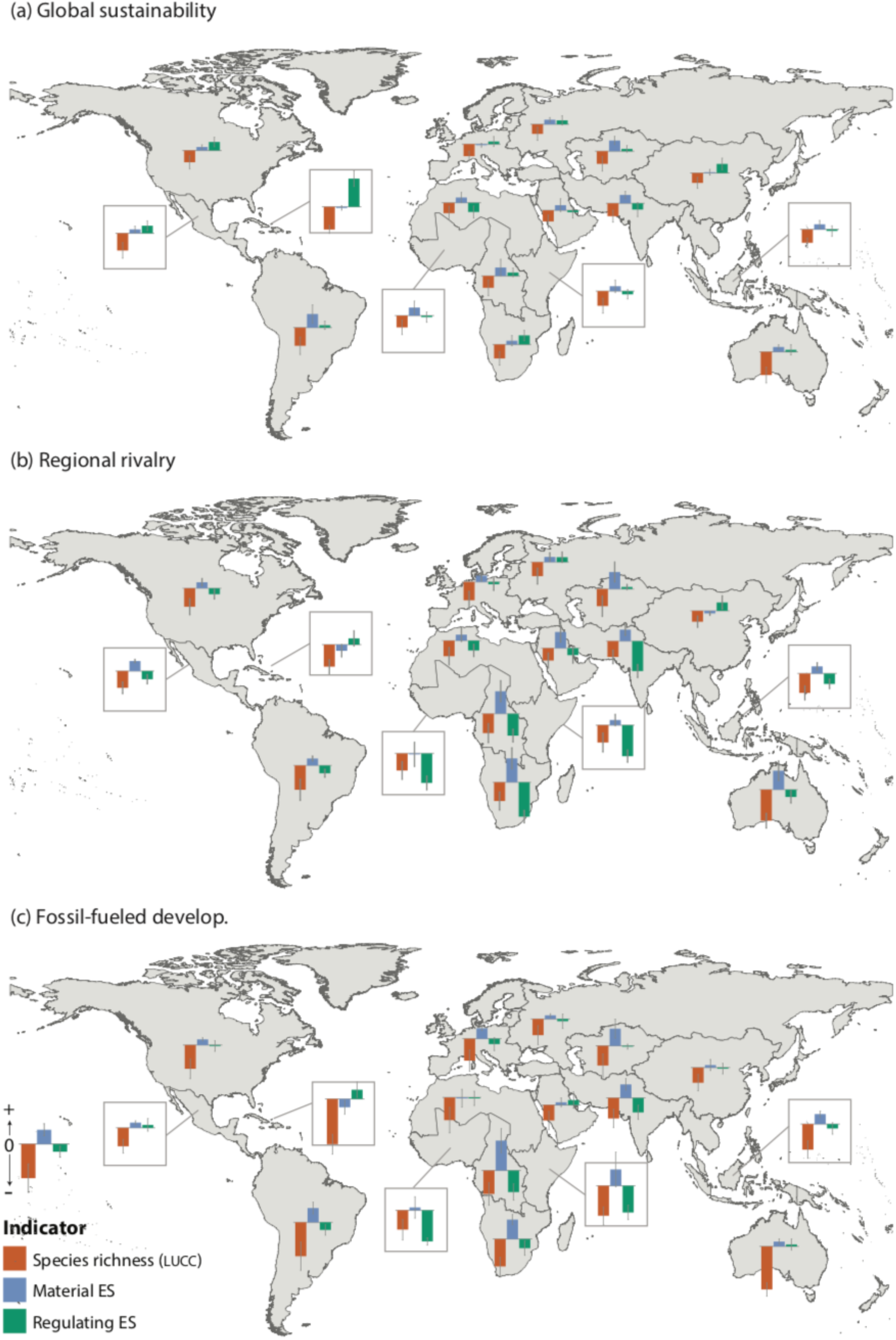
Projected regional changes in biodiversity and ecosystem services from 2015 to 2050 for (a) Global Sustainability, (b) Fossil-fueled development. Barplots show the average of the normalized values across biodiversity, material ecosystem service, and regulating ecosystem service models. Error bars are standard errors.

As with biodiversity, there is high spatial heterogeneity in future ecosystem service dynamics (Figure S9). In the fossil fueled development and regional rivalry scenarios, some regions - Central Africa, East Africa, Southern Africa, South America and South Asia – are projected to see increases of provisioning ecosystem services at the cost of substantial declines of regulating services and biodiversity (Figure 4b and 4c). Some regions such as Oceania, Mesoamerica and North Africa exhibit much lower declines in regulating services in the fossil fueled development scenario than in the regional rivalry scenario. In the global sustainability scenario, the trade-offs are smaller with some regions even registering increases in both provisioning and regulating services, such as the American regions, Eastern Europe, Southern Africa, Central Africa (Figure 4a). However, regional biodiversity still declines in most regions, as significant climate change still happens, and to a lesser extent, as a consequence of land-use change.

Our results suggest that climate change might become a more important driver of biodiversity loss than land-use change by mid-century, in agreement with recent findings based on single metrics (*14*) and in contrast to an earlier review (*10*). One reason for this finding is that future rates of land-use change are not projected to increase in any of the scenarios examined here relative to the last century rates (Figure S1a). This contrasts with two of the climate change scenarios, where rates of temperature change will still increase in the future (Figure S2). However, these results need to be interpreted with caution. There are differences in how biodiversity models capture the impacts of climate and land-use change and in the spatial grain of these impacts (*21*). Biodiversity models typically use empirical relationships at the local scale between habitat conversion and biodiversity responses and project those relationships at larger scales (*22*). In contrast, the impacts of climate are based on statistical models relating current climate with coarse species distribution patterns and assume that those relationships will hold in the future (*23*). Thus, projections for land-use change impacts are based on observed local impacts while projections for climate change are inferred from macroecological distribution patterns. In addition, our predictions assumed no species migration from climate change in any of the models, while responses to land-use change in some models allowed for species migration or species richness increases (Table S2).

Our analysis suggests that during the 20^th^ century the planet lost almost 0.8% of species from land-use change impacts alone, roughly 70,000 species if one assumes the planet’s diversity to be approximately 9 million species (*24*). This rate may vary across taxa, but is consistent with vertebrate extinctions documented by the IUCN (*2*), although some of the documented extinctions have been caused by other drivers which are not included in our models, particularly invasive alien species and direct exploitation. This agreement is even more apparent when one consider the time lags between habitat loss and extinction (*25*), which suggest that some extinctions from historical land-use change are still forthcoming. We also estimate that reductions in local species richness during the last century are around 0.9%. This contrasts with recent studies that have found no trends in local species richness in global meta-analysis of community time series (*26, 27*). Criticisms to these meta-analysis have emphasized spatial sampling biases, limited duration of time series, and the response metric used (*28*). Our analysis suggests an additional explanation: the signal may be too small to be detectable amongst the noise in available time series.

With the negotiations for a post-2020 strategy and targets underway by the parties to the Convention on Biological Diversity, our scenario analysis delivers a much-needed examination of a range of possible futures and their consequences for biodiversity and ecosystem services. Recently, it has been proposed that society must move from targets about reducing extinction rates to targets for bending upwards the curve of biodiversity loss (*29*). The global sustainability scenario comes close to achieving this for land use only, but even the modest climate change in this scenario leads to an acceleration of biodiversity loss. In addition, we see a much smaller trade-off between provisioning and regulating services in this scenario. These results provide some hope for better protection of biodiversity, particularly because the examined scenarios do not deploy all the policies that could be put in place to protect biodiversity in the coming decades. For instance, in the global sustainability scenario there is still a loss of pasture and grazing land, which are important habitats for many species, further declines in primary vegetation which is a major global driver of species extinctions (*30*), and bioenergy deployment which despite contributing to mitigate climate change can also reduce species habitats (*31*). Introducing further measures such as further regulation of deforestation, increasing effectiveness of protected areas (*32*), stronger changes in consumption patterns (*33*), and sensible natural climate solutions (*34*), could result in even better prospects for biodiversity and ecosystem services. We need to develop a novel generation of global scenarios that aim at achieving positive futures for biodiversity (*35*), to identify better development policies and biodiversity management practices.

## Acknowledgments

This study was carried out with the support of the IPBES Expert Group on Scenarios and Models and its Technical Support Unit.

## Funding

HP, IR, IM, HK, JS and NT were supported by the German Research Foundation (DFG FZT 118) which also provided funding for a workshop.

## Author contributions

RA, PL, HP, DV, AP and GH conceptualized the work. Analysis, curation and visualization were carried out by IR, IM and HK. All authors contributed models and data or ideas for the analysis. HP wrote the paper with contributions from all authors.

## Competing interests

Authors declare no competing interests; and

## Data and materials availability

All data and code used in the analysis are available from the authors upon request.

## Supplementary Materials

Materials and Methods

Figures S1-S9

Tables S1-S3

References (*36-80*)

## Notes

### Competing Interest Statement

The authors have declared no competing interest.

### Summary of Updates

Author order in biorXiv was revised to match author order in manuscript.

