## Supplementary Materials for "Global trends in biodiversity and ecosystem services from 1900 to 2050"

**This PDF file includes:**

Materials and Methods
Figs. S1 to S9
Tables S1 to S3

**Materials and Methods**

This study was conducted under the auspices of the Expert Group on Scenarios and Models of the Intergovernmental Science-Policy Platform on Biodiversity and Ecosystem Services (IPBES). The detailed protocol of this multi-model study was published in (*17*). Below we summarize the main methodological aspects.

Scenarios

All models used the same set of scenarios: SSP1 with RCP2.6 ("global sustainability" with low land-use pressure and low level of climate change, (*36*)), SSP3 with RCP6.0 ("regional rivalry" with high land-use pressure and intermediate level of climate change, (*37*)), and SSP5 with RCP8.5 ("fossil-fueled development" with intermediate land-use pressure and high level of climate change, (*38*)) – to assess a broad range of plausible futures (Table S1). We used land-use projections for these scenarios ignoring the impacts of climate change, although the deployment of land-based climate mitigation strategies is considered in connection to each of the SSP-RCP combinations. Land-use projections for SSP3xRCP6.0 were not available, so we chose the closest land-use projections available, SSP3xRCP7.0.

Land use data

All models used the Land Use Harmonization (*39–43*) version 2 dataset (LUH2, see http://luh.umd.edu/data.shtml for data). LUH2 provides global gridded land-use datasets at 0.25° resolution with annual time-steps comprising estimates of historical land-use change (850-2015) and future projections (2015-2100) under the assumptions of each Shared Socio-economic Pathway (SSP) (*44*). The 12 land use categories (Table S3) include the separation of primary and secondary natural vegetation into forest and non-forest sub-types, pasture into managed pasture and rangeland, and cropland into multiple crop functional types (C3 annual, C3 perennial, C4 annual, C4 perennial, and C3 nitrogen-fixing crops). The LUH2 dataset also computes all transitions between these 12 land use types, resulting in over 100 possible transitions per grid cell per year (e.g., crop rotations, shifting cultivation, agricultural changes, wood harvest) as well as various agricultural management layers (e.g., irrigation, synthetic nitrogen fertilizer, biofuel crops). Due to specific model parameterizations, each biodiversity and ecosystem service model used its own aggregation of the land use categories (see (*17*) for more details).

Climate data

Models used historical climate data and future projections associated with each SSPxRCP combination (*20*) from CMIP5 / ISIMIP2a (*45*) or its downscaled version from the WorldClim (*46*), or the projections from MAGICC 6.0 (*47*, *48*). Most models used the IPSL-CM5A-LR (*49*) projections which are mid-range across the 5 GCMs in ISIMIP2a (*50*) – that includes 12 climate variables at 0.5° resolution on daily time steps from the pre-industrial period 1951 to 2099 (*45*). The WorldClim downscaled dataset has 19 bioclimatic variables monthly from 1960 to 1990 and multi-year averages for specific points in time (e.g., 2050, 2070) up to 2070 at 1km resolution. MAGICC 6.0 climate data (*47*, *48*) in the IMAGE model framework (*51*) was used for the GLOBIO model.

Biodiversity models

All models have been published in peer-reviewed journals, although in some cases modifications have been made to the original model (see (*17*) for details in modifications). In total, 8 spatially-explicit models were used (Table S2), these include three species distributions models - AIM-biodiversity (*52*), InSiGHTS (*53*, *54*), MOL (*55*, *56*); and five community models (cSAR-iDiv (*57*), cSAR-IIASA-ETH (*58*), BILBI (*59*), PREDICTS (*60*, *61*), GLOBIO (*62*, *63*). Three of these models, BILBI, PREDICTS and cSAR-iDiv share coefficients for the impacts of land-use on biodiversity from the PREDICTS database (*61*). The biodiversity models have different methodological approaches, taxonomic groups, spatial resolution and output metrics (Table S2), but they were harmonized as described below.

<u>Ecosystem services models</u>

For ecosystem functioning and services, five spatially-explicit models were used. They include three process-based DGVM models – LPJ-GUESS (*64–66*), LPJ (*67*, *68*), and CABLE-POP (*69*) – and two ecosystem services models – InVEST (*70*) and GLOBIO-ES (*71*, *72*)). These rely on different modelling approaches to estimate a wide range of biophysical outputs, which were harmonized as described in the next sections (see Table S2 for a summary of the models, details available in (*17*)).

<u>Scales of analysis (local, regional and global) and harmonization of metrics</u>

Model outputs were produced at three spatial scales: one-degree grid cells ($\alpha$ metrics), at the regional level (regional $\gamma$ metrics) for the 17 IPBES sub-regions (*73*), and at the global level (global $\gamma$ metrics). The methodology adopted by each modelling team to aggregate from the original resolution of the model to one-degree cells was the arithmetic average of the values in the original resolution.

The model outputs addressed very different facets of biodiversity (e.g., species ranges, local species richness, global species extinctions, abundance-based intactness, and compositional similarity), as well as different facets of ecosystem services (e.g., pollination, carbon sequestration, soil erosion, wood production, nutrient export, coastal vulnerability), often with little overlap between different models. In addition, even for the same facet of biodiversity or ecosystem service, different models outputted different metrics. In order to ensure comparability, output metrics for each model were converted to proportional changes relative to the beginning time of the analysis (e.g., $\Delta y = \frac{y_{t1} - y_{t0}}{y_{t0}}$), where $y_t$ is the value of the metric at time *t*, and $t_0$ and $t_1$ are respectively the beginning and the end of the time period. In addition, models that simulated a continuous time series of climate change impacts calculate $y_t$ as 20-year averages around the midpoint *t* in order to account for inter-annual variability.

<u>Biodiversity metrics</u>

Outputs of each biodiversity model were assigned to one or more of the following harmonized biodiversity metrics (Table S2): species richness (S), mean species habitat extent ($\dot{H}$), and species-abundance based biodiversity intactness (I). While all metrics were reported as proportional changes relative to the beginning of a time period, intactness was also reported as a score relative to a pristine baseline. For mapping purposes, local changes in proportional species richness were converted in normalized changes in absolute species richness ($\Delta SS$), by multiplying by the number of species in each cell divided by the number of species in the richest cell. Global spatial averages of the local metrics were calculated across all terrestrial one-degree cells and are denoted with an overbar (e.g. $\overline{\Delta S_\alpha}$) to distinguish it from averages of a metric across species ($\dot{H}$).

In the end, the harmonized metrics analyzed were:

- $\Delta S_\alpha(x, y) = \frac{S_\alpha(x,y,t1) - S_\alpha(x,y,t0)}{S_\alpha(x,y,t0)}$, where $S_\alpha(x, y, t)$ is the number of species at cell (x,y) at time *t*;

- $\Delta SS_\alpha(x,y) = \Delta S_\alpha(x,y) \times \frac{S(x,y)}{Max_{\{x,y\}}[S(x,y)]}$, where $S(x,y)$ is the number of species at cell (x,y) calculated from current species distribution maps, and the maximum value is calculated across all cells;
- $\Delta S_\gamma(region) = \frac{S_\gamma(region,t1) - S_\gamma(region,t0)}{S_\gamma(region,t0)}$, where $S_\gamma(region,t)$ is the number of species in an IPBES sub-region or in the globe at time t;
- $\Delta \dot{H}_\gamma = \frac{1}{S\gamma}\sum_{i=1}^{S\gamma}\frac{H_\gamma(i,t1,i) - H_\gamma(i,t0)}{H_\gamma(i,t0)}$, where $H_\gamma(i,t)$ is the global habitat extent of species *i* at time *t*;
- $I_\alpha(x,y,t)$, which is the species-abundance based intactness value for cell (x,y) at time *t* relative to a pristine baseline, with 100% corresponding to a pristine habitat and 0% to a completely degraded habitat.

In addition, global spatial averages for $\alpha$ metrics where calculated as follows:

- $\overline{\Delta S_\alpha} = \sum_{x,y}\frac{\Delta S_\alpha(x,y)}{n}$
- $\overline{\Delta \dot{H}_\alpha} = \sum_{x,y}\frac{\Delta \dot{H}_\alpha(x,y)}{n}$
- $\overline{I_\alpha} = \sum_{x,y}\frac{I_\alpha(x,y)}{n}$

where *n* is the number of terrestrial one-degree cells.

The harmonized biodiversity metrics need to be interpreted with care as the original model outputs mapped to the same harmonized metric can differ in some technical details. For instance, the GLOBIO model (*62*, *63*) outputs a metric called "Mean Species Abundance" (MSA) that represents "the mean abundance of original species in relation to a particular pressure as compared to the mean abundance in an undisturbed reference situation"; likewise the PREDICTS model (*74*) outputs a metric called "Biodiversity Intactness Index (BII)" that represents "the average abundance of originally present species across a broad range of species, relative to abundance in an undisturbed habitat". While both metrics have been harmonized as representing species-abundance based intactness (*I*), they are calculated differently in the models (i.e., the former is the average of abundance ratios while the latter is the ratio of the sums). Similarly, models based on the species-area relationship (*75*) produced similar metrics (relative change in species richness) but covered different taxonomic groups (Table S2).

Ecosystem services metrics

A similar effort was made to assign the metrics outputted by the ecosystem function and services models to a set of harmonized metrics (Table S1). We used the typology of the IPBES Nature's Contributions to People (NCPs) (*19*) to classify material and regulating services. For each of the following ecosystem services we assigned one biophysical metric from one or more models, sometimes changing the sign of the reported metric for consistency: bioenergy production; food and feed production; timber production; ecosystem carbon; crop pest control (more is better control); coastal resilience (more is greater resilience); pollination; soil protection; nitrogen retention (more is higher water quality).

The dynamic global vegetation models (DGVMs) tend to output similar metrics and have similar assumptions (*76*), but the two ecosystem service models (GLOBIO and InVEST) tended to output different metrics for the same service. DGVMs have been used in the climate change modeling community for decades so they benefit from a long history of multi-model inter-comparison (*77*). Therefore, while for certain metrics, such as ecosystem carbon pool, the metrics are calculated in a similar way and use equivalent biophysical units (e.g. Kg C), for other metrics, e.g., pollination, direct comparison of absolute values was not feasible. For instance, GLOBIO-ES (*72*, *78*) defines their metric of pollination services as "the fraction of cropland potentially pollinated, relative to all available cropland", but in InVEST (*79*) defines it as "the proportion of agricultural lands whose pollination needs are met". As for biodiversity metrics, this problem was addressed by using proportional changes of each metric in each model at each scale of analysis.

Comparison of biodiversity, regulating and material ecosystems services

To understand how biodiversity and ecosystem services varied concurrently in each IPBES sub-region (Figure 4) we mapped regional changes in biodiversity and in aggregated regulating and material ecosystem services, from 2015 to 2050 for all three scenarios. First, we normalized changes in regional species richness ($\Delta S_{\gamma}$) and ecosystem service metrics for all scenarios and regions, by dividing the proportional changes for each sub-region and scenario and model metric by the maximum value of that metric for all subregions in all scenarios. In this way, we obtained a normalized $\Delta Y$ with values between -1 and +1 for biodiversity or ecosystem service metric in each region and scenario. Next, we clustered all normalized model values into biodiversity metrics, material ecosystem services and regulating ecosystem services.

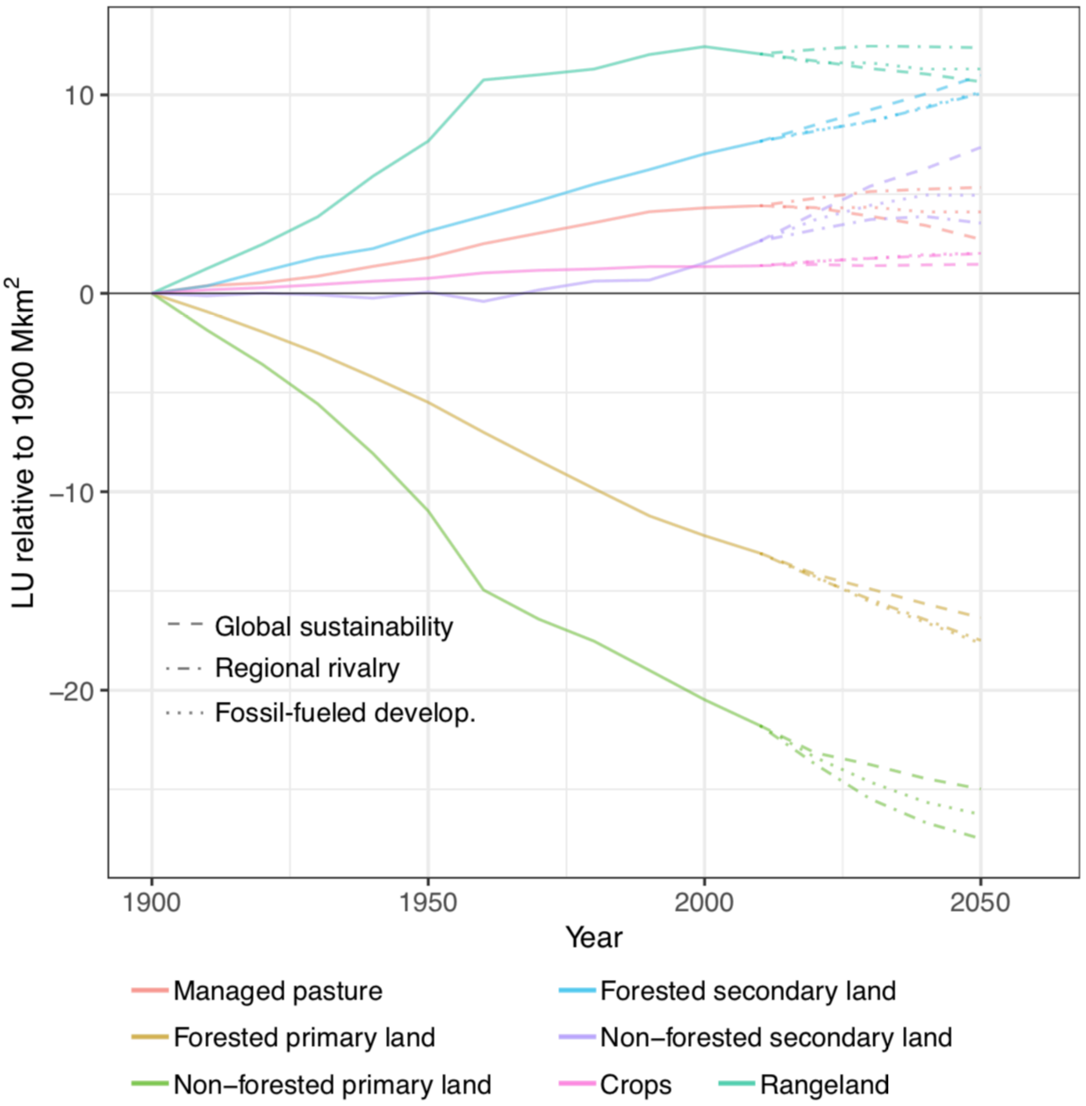


**Fig. S1. (a)** Global historical trends (1990-2015) in land-use and projected trends for each scenario (2015-2050). Lines correspond to absolute area changes relative to the year 1900. The original area covered by each land-use in 1900 was: forested primary land (36.0 $Mkm^2$), non-forested primary land (50.7 $Mkm^2$), forested secondary land (6.3 $Mkm^2$), non-forest secondary land (11.8 $Mkm^2$), managed pasture (3.5 $Mkm^2$), rangeland (12.9 $Mkm^2$), cropland (9.5 $Mkm^2$).

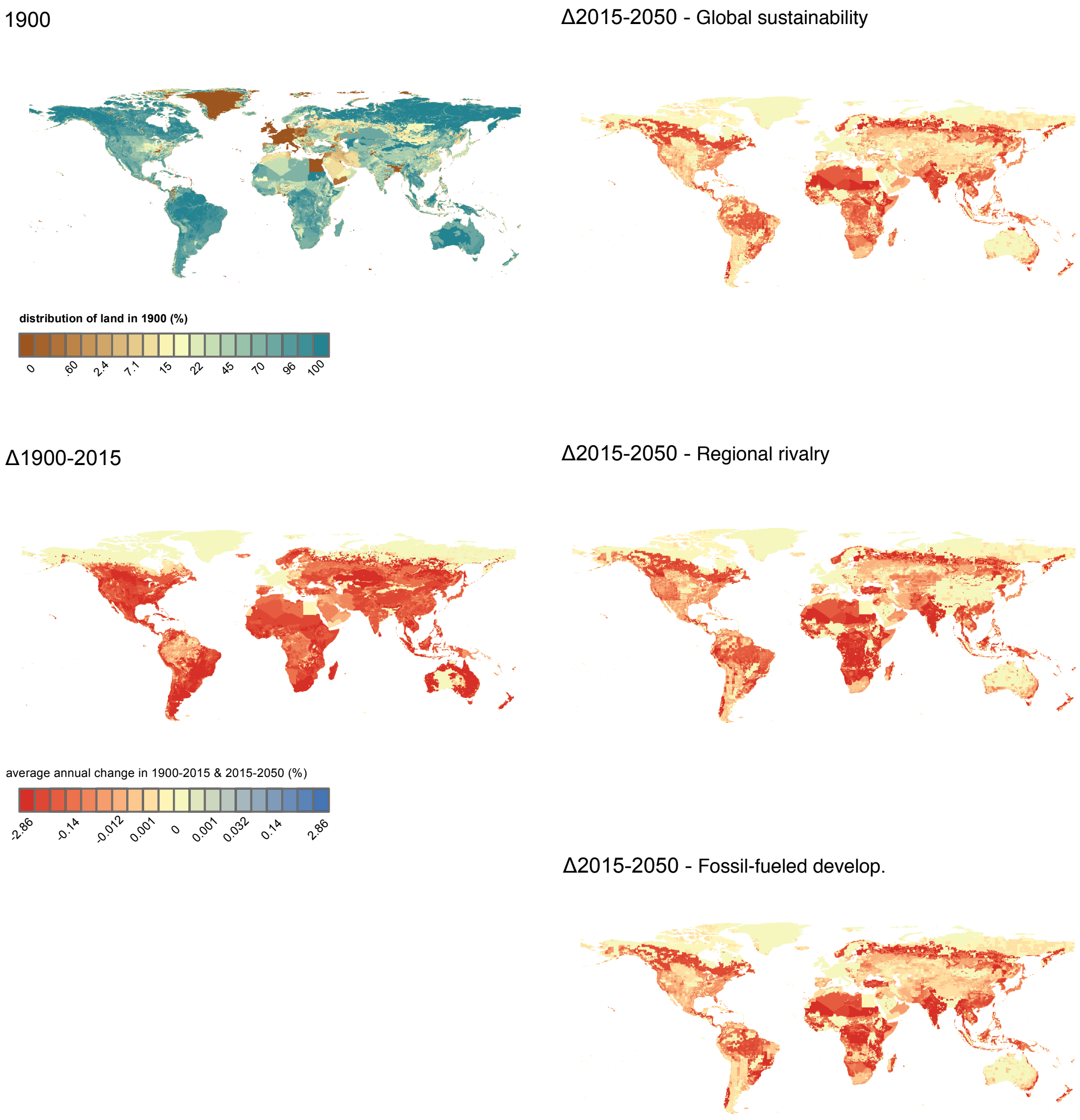


**Figure S1 (b)** Distribution of primary land (forest & non-forest) in 1900, historical changes (1900-2015) and future changes (2015-2050) in each scenario. Please note that changes are reported in absolute percentage points (i.e., $y_{t1}$-$y_{t0}$ where y is the percentage of the area in a cell covered by that land use type). Color scales are based on quantile intervals considering all land cluster types for i) 1900 and ii) the past (Δ1900-2015) and future (Δ2015-2050) combined.

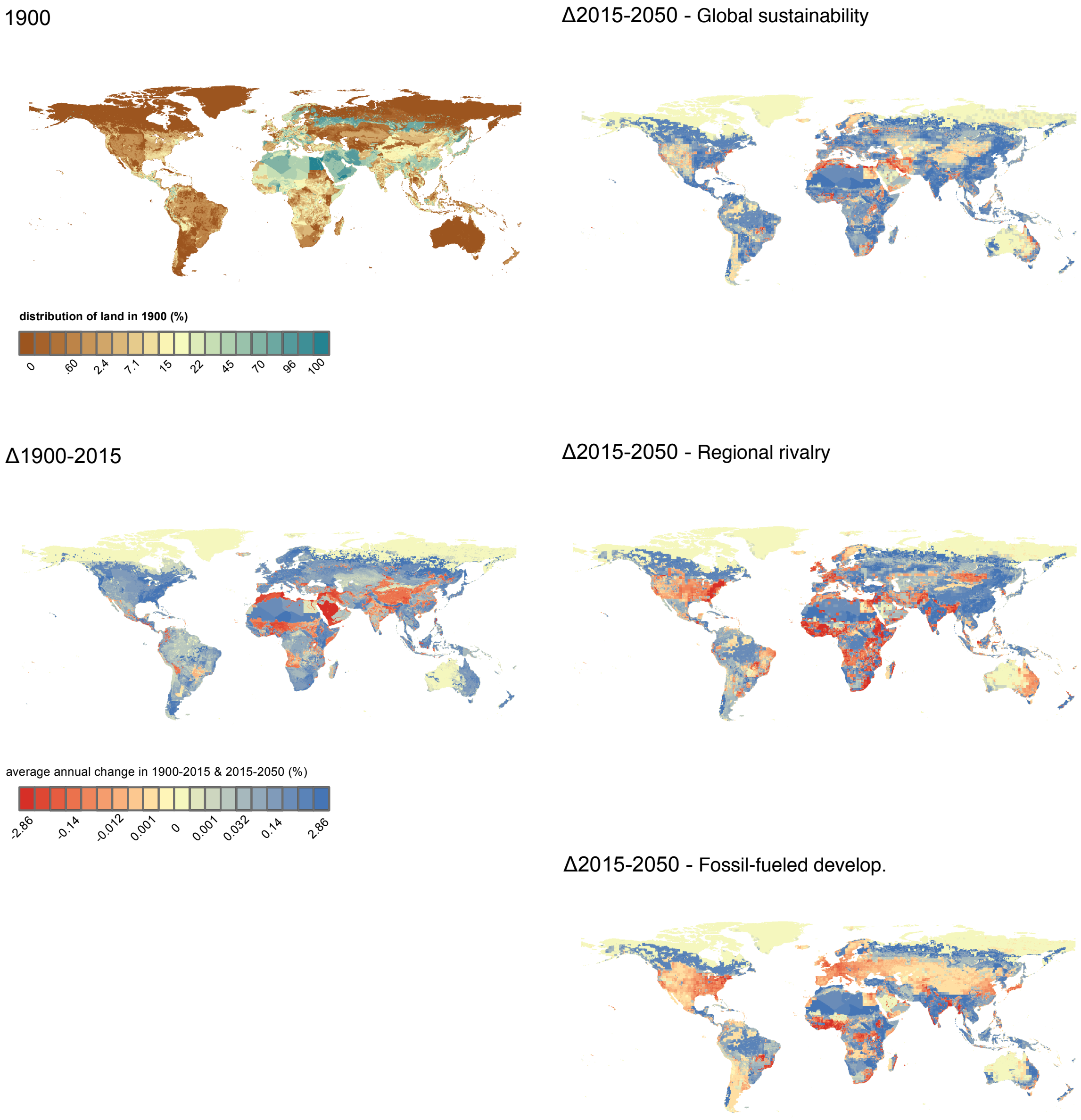


**Figure S1 (c)** Distribution of secondary land (forest & non-forest) in 1900, historical changes (1900-2015) and future changes (2015-2050) in each scenario. Please note that changes are reported in absolute percentage points (i.e. $y_{t1}$-$y_{t0}$ where y is the percentage of the area in a cell covered by that land use type). Color scales are based on quantile intervals considering all land cluster types for i) 1900 and ii) the past (Δ1900-2015) and future (Δ2015-2050) combined.

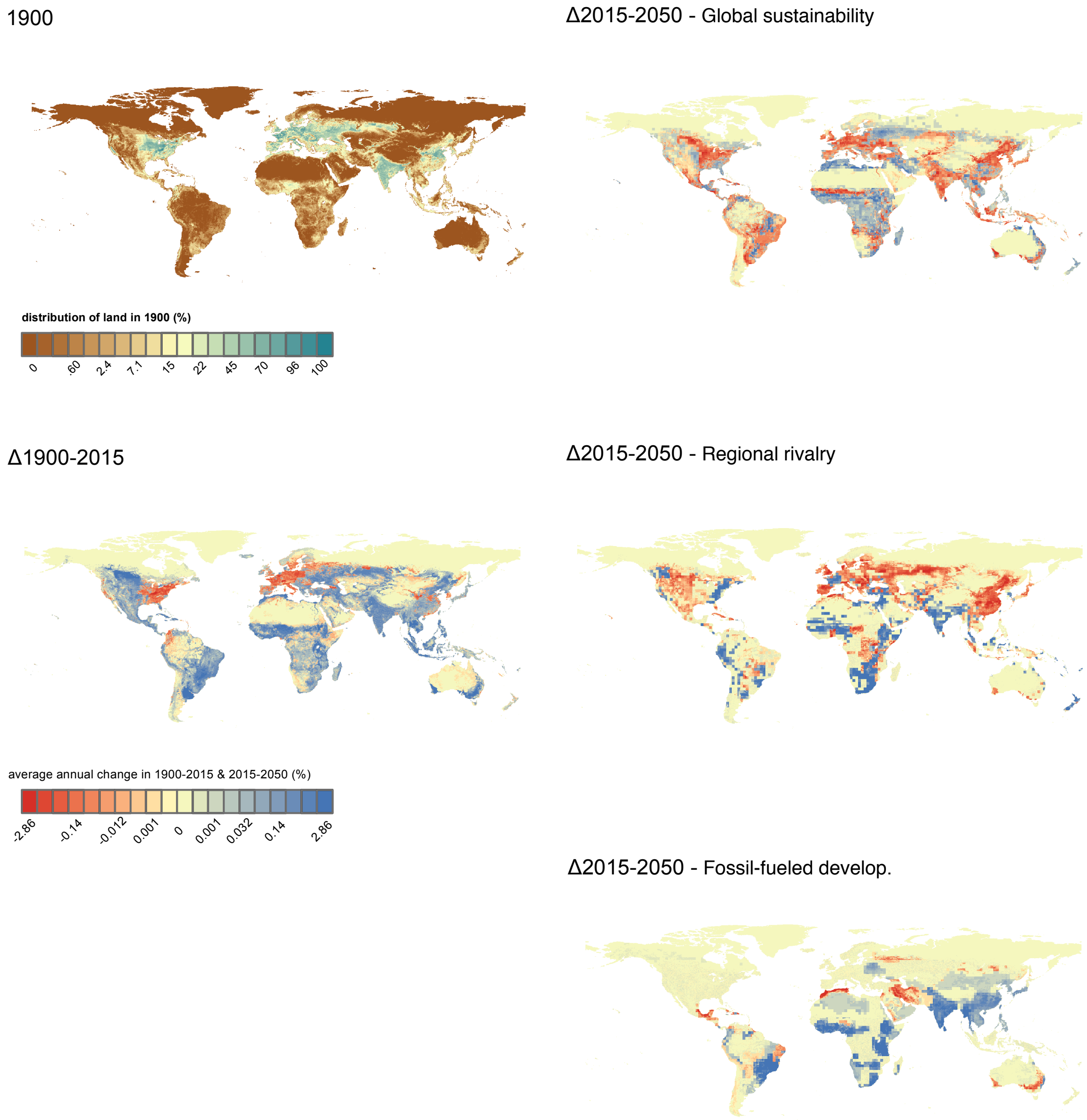


**Figure S1 (d)** Distribution of cropland (C3 & C4) in 1900, historical changes (1900-2015) and future changes (2015-2050) in each scenario, in percentage. Please note that changes are reported in absolute percentage points (i.e. $y_{t1}$-$y_{t0}$ where y is the percentage of the area in a cell covered by that land use type). Color scales are based on quantile intervals considering all land cluster types for i) 1900 and ii) the past (Δ1900-2015) and future (Δ2015-2050) combined.

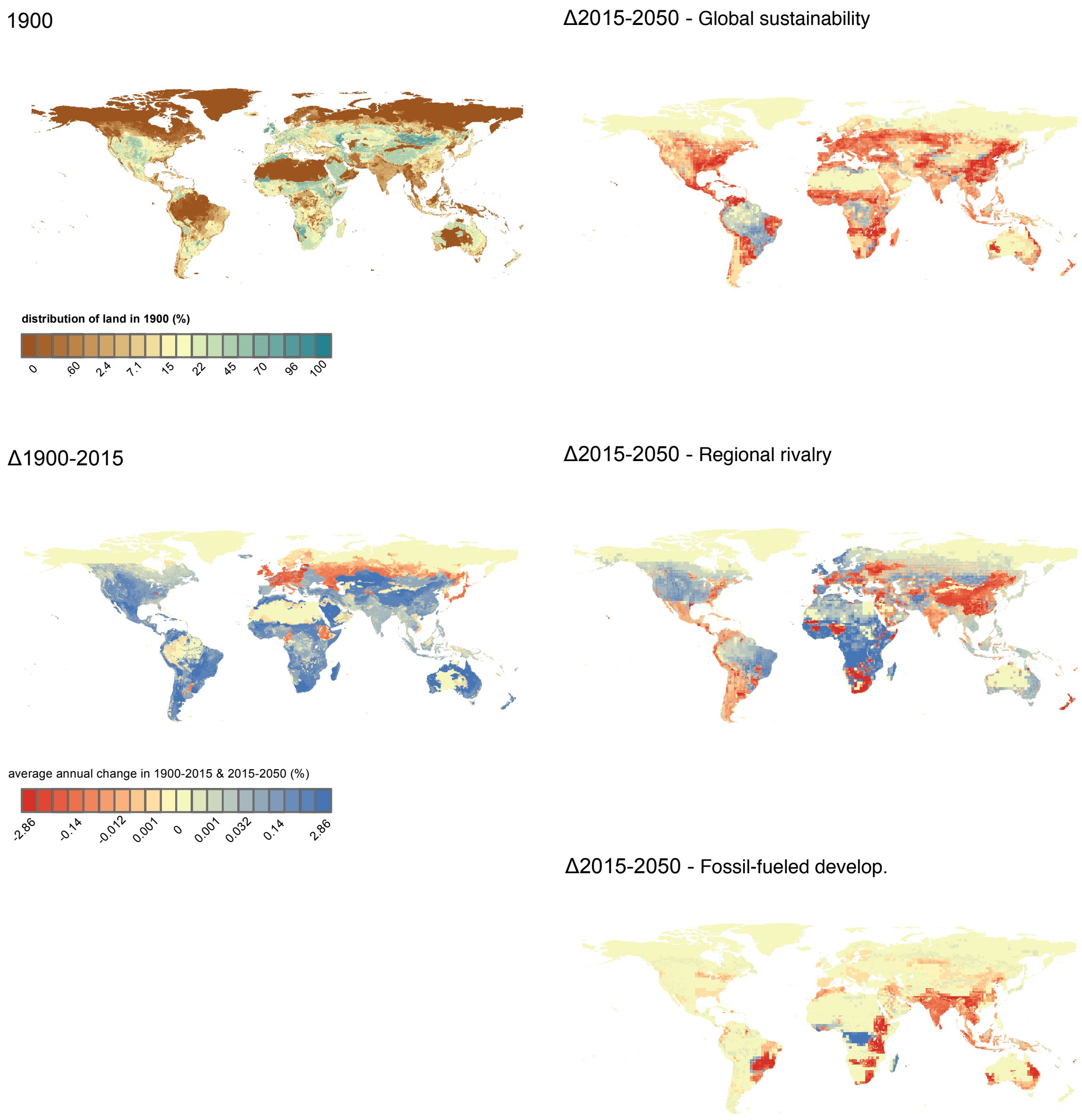


**Figure S1 (e)** Distribution of pasture and rangeland in 1900, historical changes (1900-2015) and future changes (2015-2050) in each scenario, in percentage. Please note that changes are reported in absolute percentage points (i.e. $y_{t1}$-$y_{t0}$ where y is the percentage of the area in a cell covered by that land use type). Color scales are based on quantile intervals considering all land cluster types for i) 1900 and ii) the past (Δ1900-2015) and future (Δ2015-2050) combined.

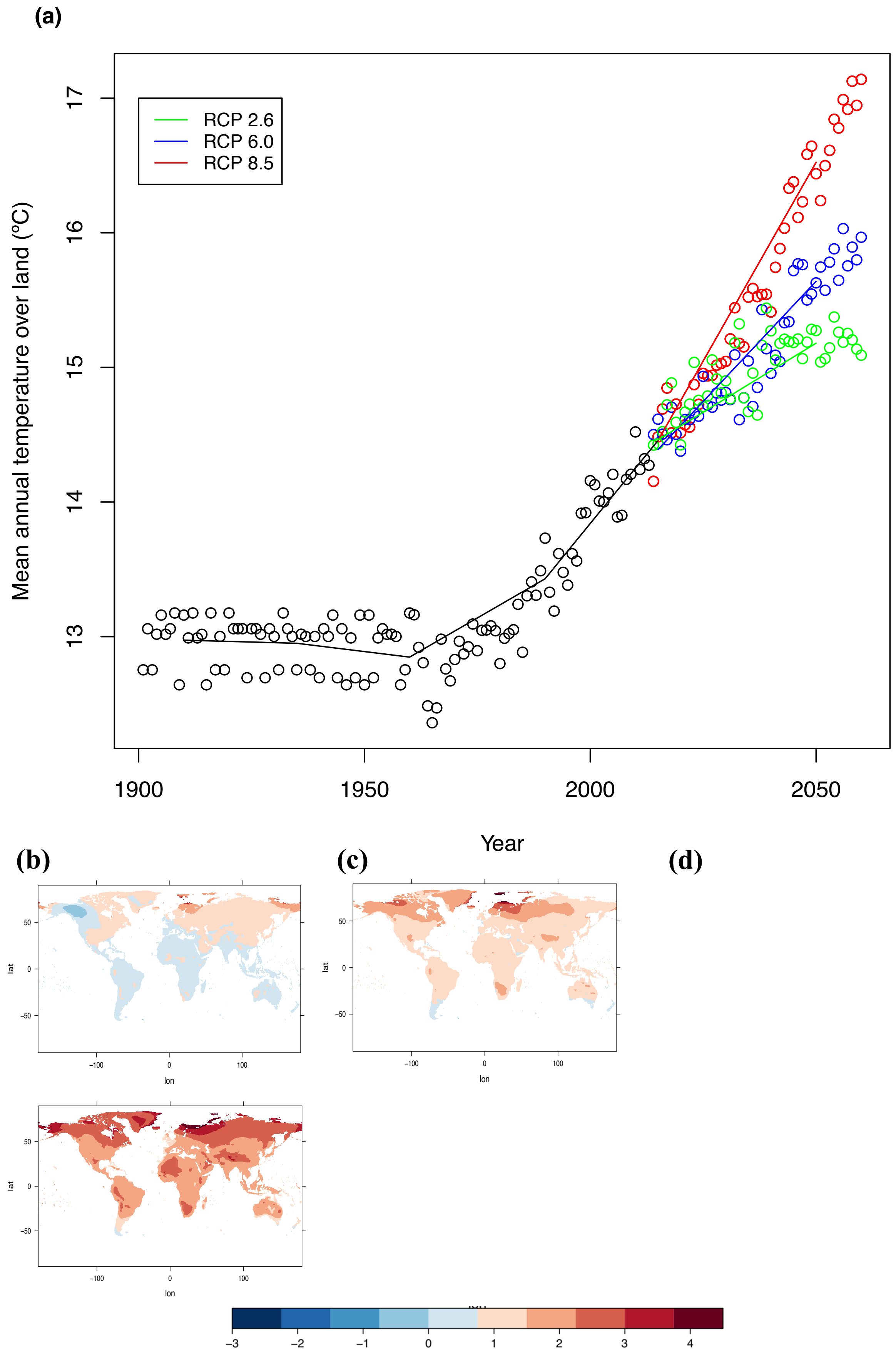


**Fig. S2. (a)** Global historical trends (1990-2015) in mean annual temperature and for each scenario (2015-2050). Spatial distribution absolute changes in mean annual temperature in each scenario (2015-2050): **(b)** global sustainability - RCP2.6, **(c)** regional rivalry - RCP6.0, **(d)** fossil-fueled development - RCP8.5.

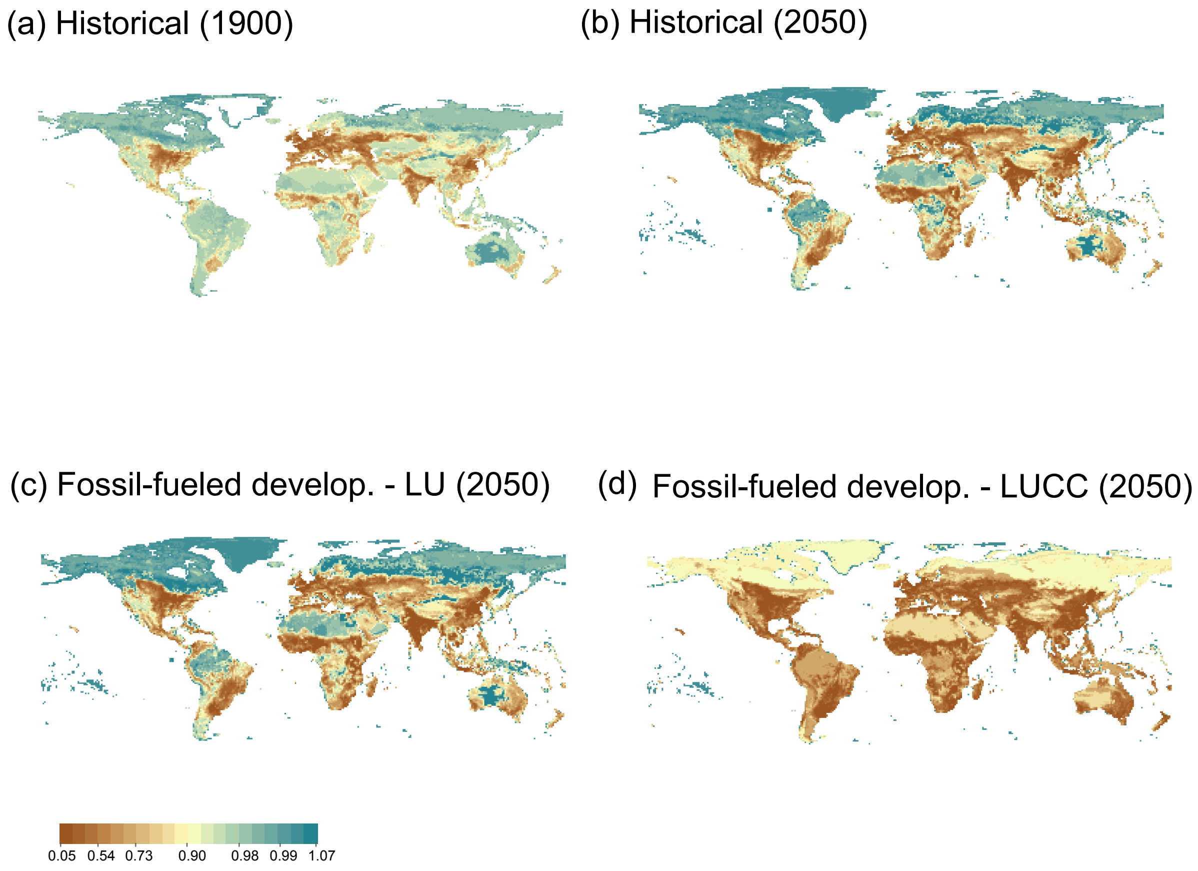


**Fig. S3.** Spatial distribution of intactness (*I*): **(a)** year 1900; **(b)** 2015; **(c-d)** 2050 in the fossil-fueled development scenario based on land-use change alone **(c)** and on the combined impacts of land-use change and climate **(d)**. Values correspond to the inter-model mean between PREDICTS and GLOBIO, except for **(d)** which is based only on GLOBIO. Values are scores relative to a pristine baseline (a score of 1 corresponds to pristine, while a score of 0 corresponds to fully degraded). Color scale is based on quantile intervals when considering all maps features.

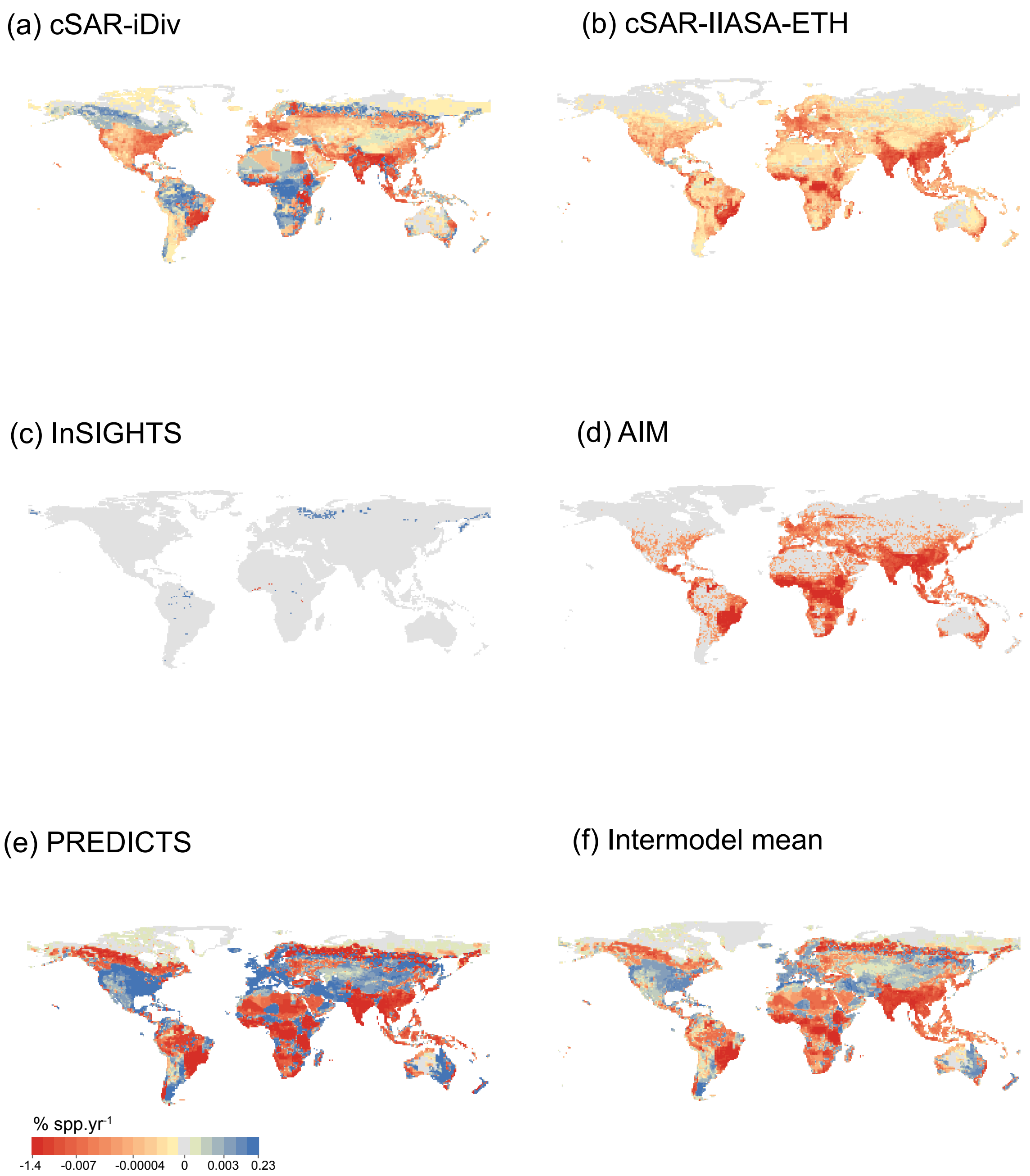


**Fig. S4.** Spatial agreement between biodiversity models. Projection of normalized changes in local species richness per year ($\Delta SS_{\alpha}$) during 2015-2050 caused by land-use change alone for the regional rivalry scenario: **(a)** cSAR-iDiv model; **(b)** cSAR-IIASA-ETH model; **(c)** InSIGHTS model; **(d)** AIM-B model; **(e)** PREDICTS model; **(f)** inter-model mean. A value of 1% yr$^{-1}$ corresponds to a decline in the number of local species equal to 1% species of the most speciose grid cell. Color scale is based on quantile intervals when considering all maps features.

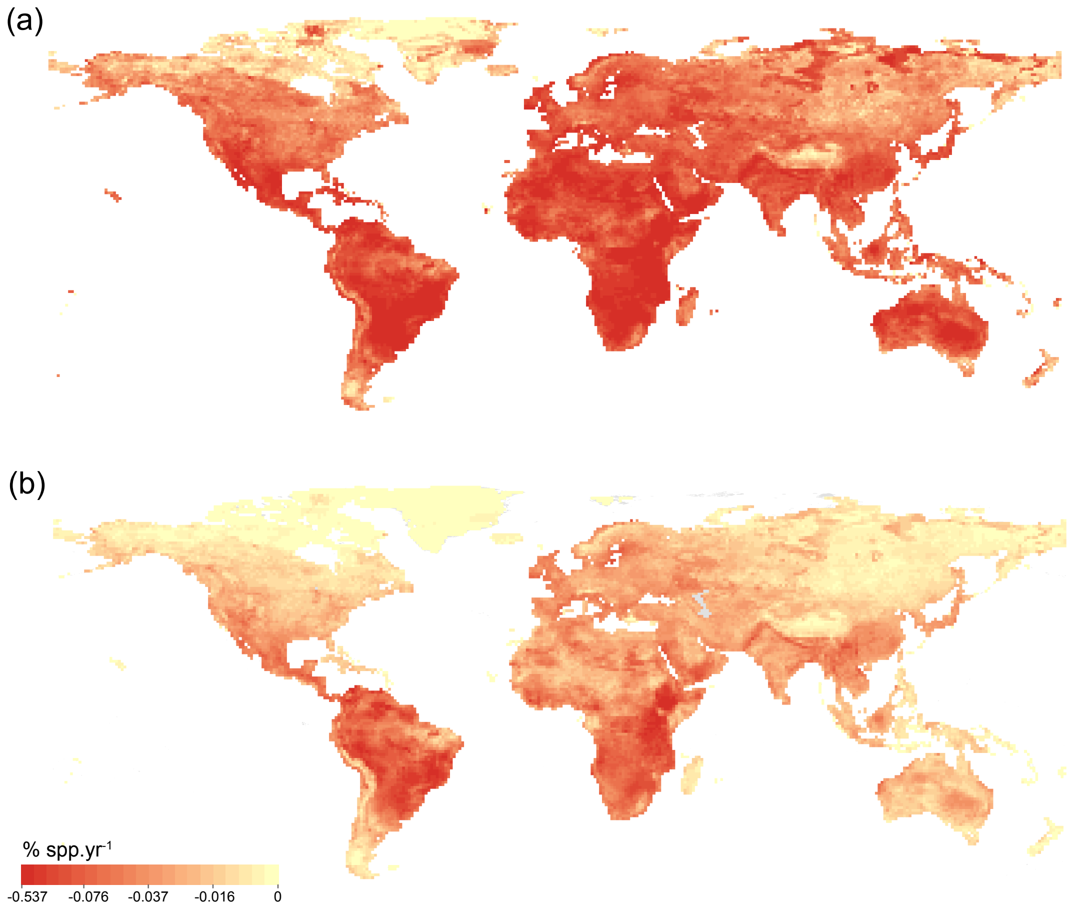


**Fig. S5.** Biodiversity metrics of the AIM model for the fossil fueled development scenario for 2015-2050: **(a)** proportional changes in local species richness ($\Delta S_{\alpha}$); **(b)** normalized changes in local species richness per year ($\Delta SS_{\alpha}$). Color scale is based on quantile intervals when considering all maps features.

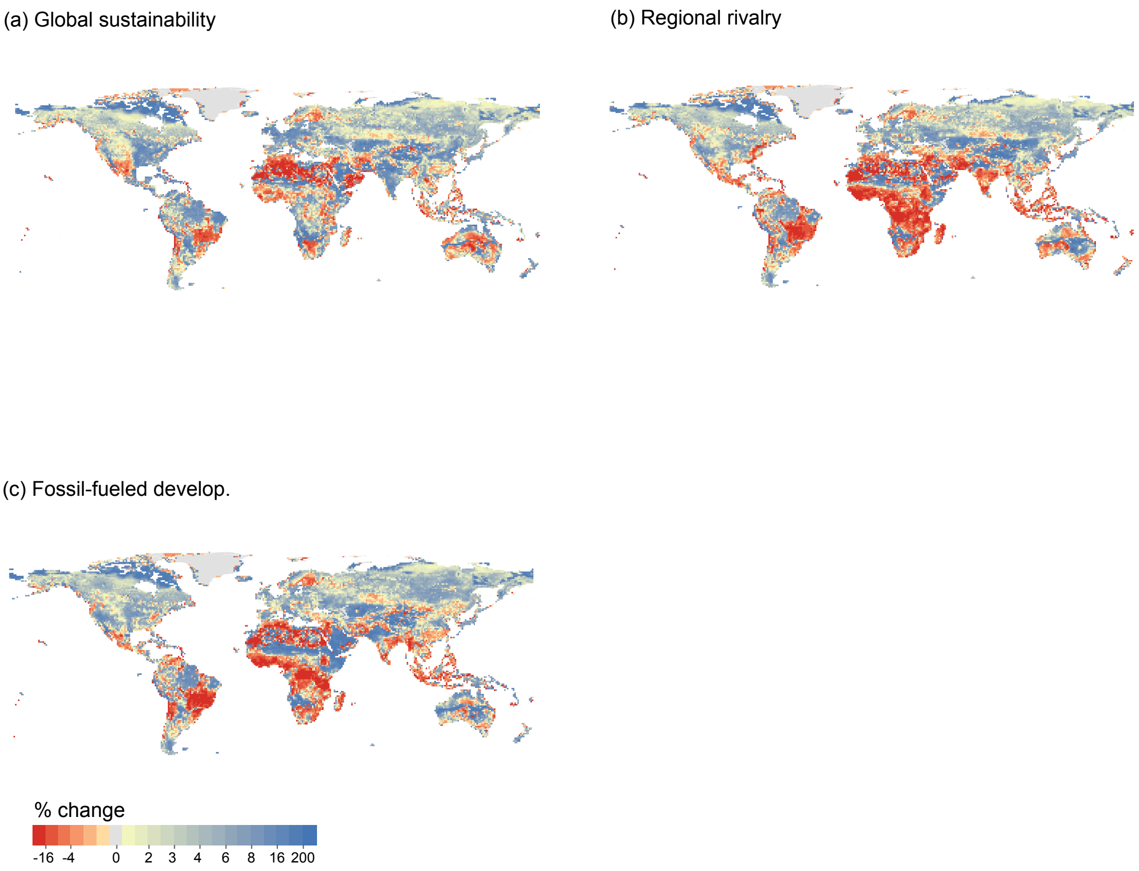


**Fig. S6.** Ecosystem carbon pools across scenarios. Inter-model mean of proportional changes for 2015-2050 (N=4, CABLE-POP, LPJ, LPJ-GUESS, GLOBIO-ES): (a) global sustainability, (b) regional rivarly, (c) fossil-fueled development. Color scale is based on quantile intervals when considering all maps features.

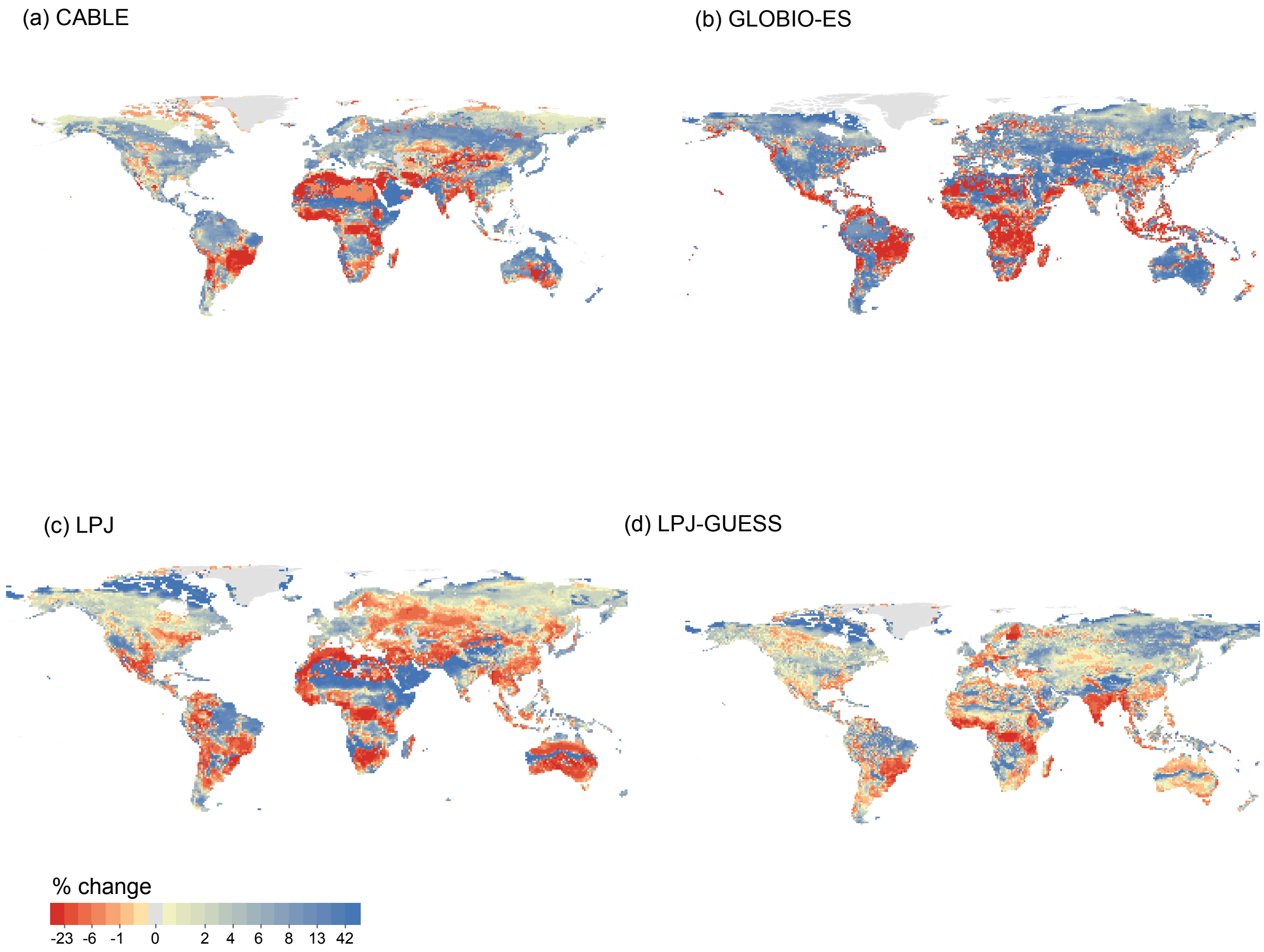


**Fig. S7.** Spatial agreement across models in ecosystem carbon for the fossil fuel development scenario for 2015-2050: **(a)** CABLE-POP, **(b)** GLOBIO-ES, **(c)** LPJ and **(d)** LPJ-GUESS. The inter-model mean can be found in Figure S7. Color scale is based on quantile intervals when considering all maps features.

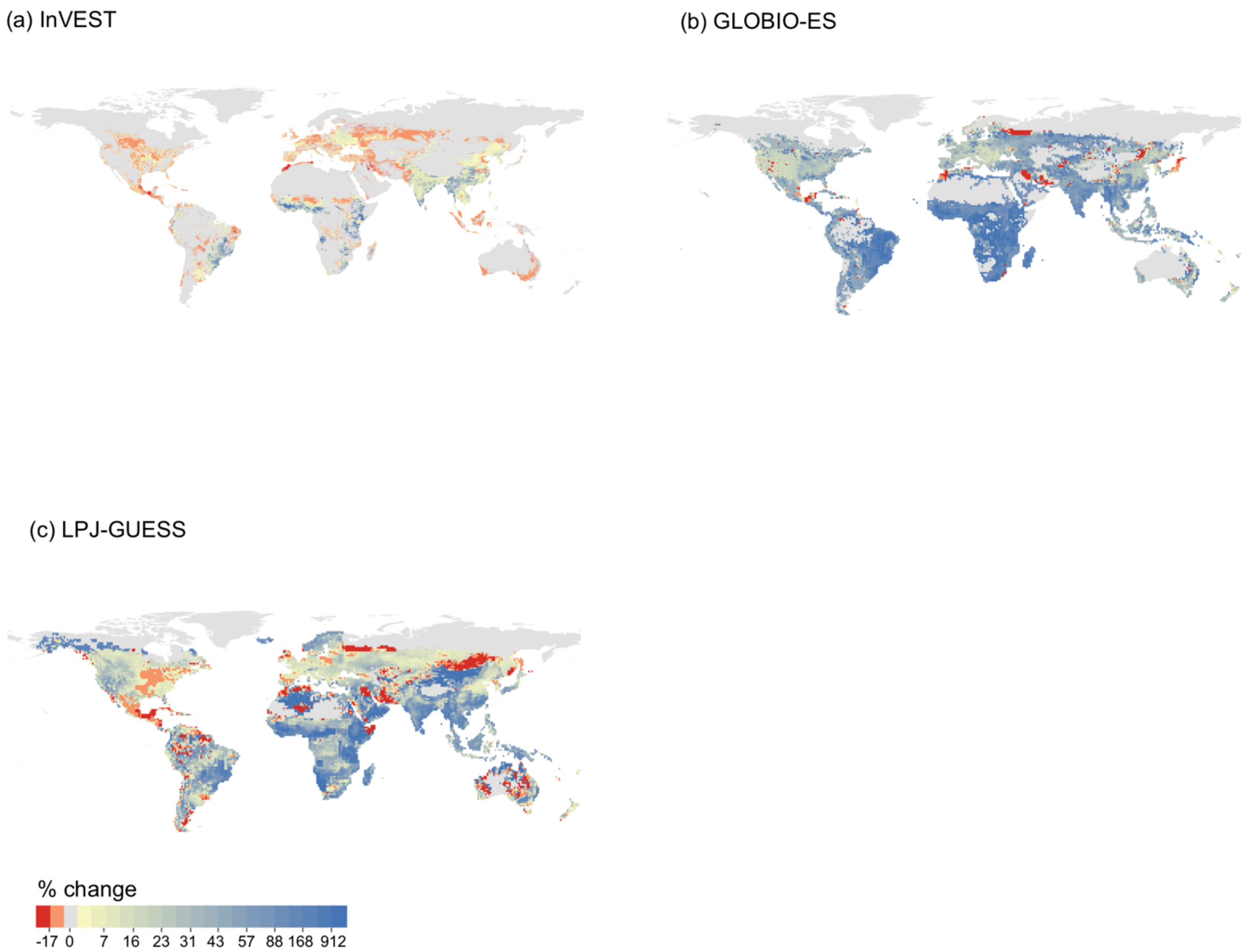


**Fig. S8.** Spatial agreement across models in modelled food and feed production for the fossil fueled development scenario for 2015-2050: **(a)** InVEST, **(b)** GLOBIO-ES and **(c)** LPJ-GUESS. The inter-model mean can be found in Figure S7. Color scale is based on quantile intervals when considering all maps features.

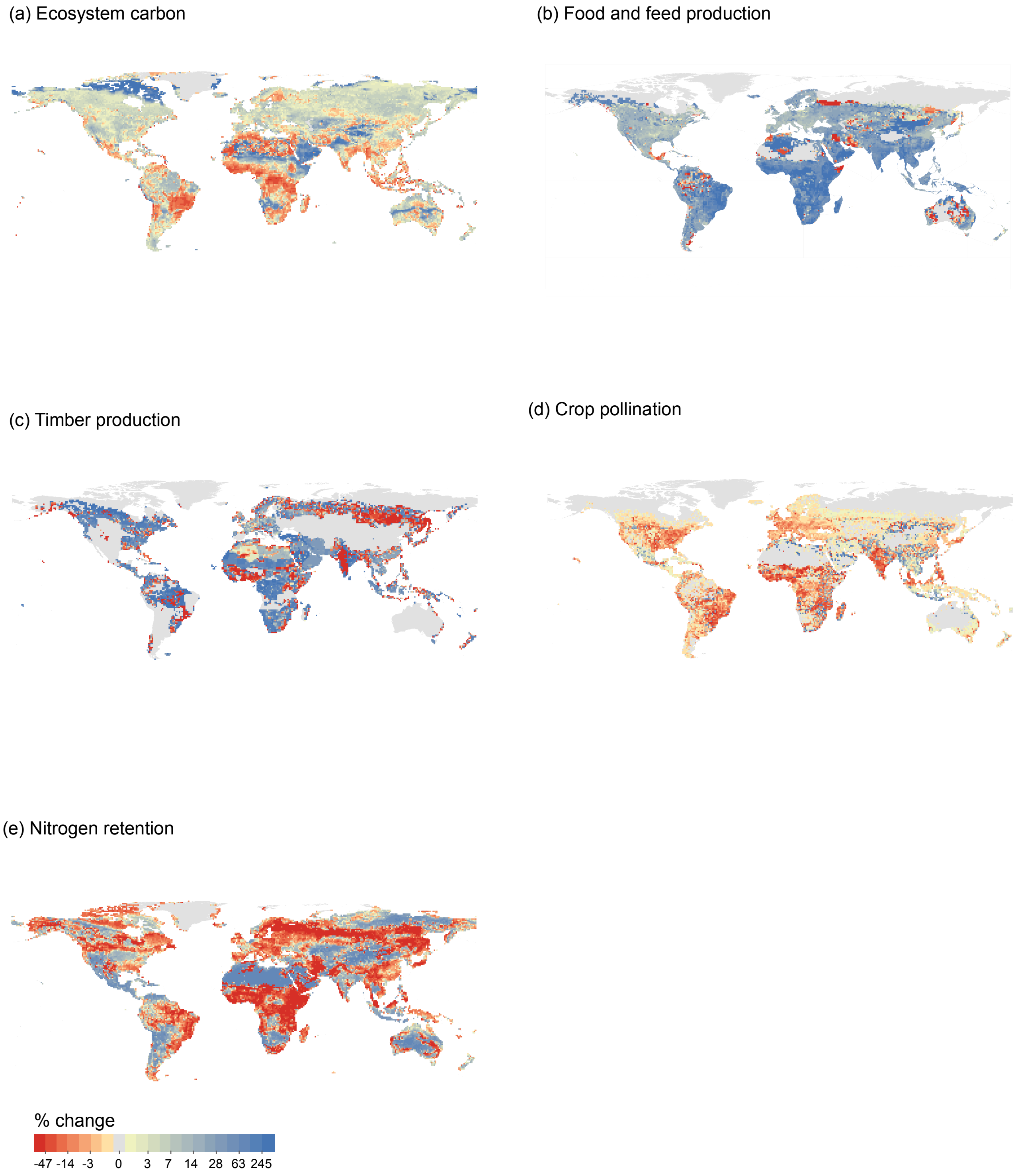


**Fig. S9.** Spatial distribution of ecosystem service changes. Inter-model mean projection of proportional changes (2015-2050) in the fossil fueled development scenario for: (a) Ecosystem carbon (N=4), (b) Food and feed production (N=3), (c) Timber production (N=2), (d) Crop pollination (N=2) and (e) Nitrogen retention (N=2). Colour scale is based on quantile intervals when considering all maps features.

**Table S1.** Characteristics of SSP and RCP scenarios (based on (*18*) and https://secure.iiasa.ac.at/web-apps/ene/SspDb/dsd?Action=htmlpage&page=about)

| | SSP1xRCP2.6<br>Global sustainability | SSP3xRCP6.0<br>Regional Rivalry | SSP5<br>Fossil-fueled Development |
|---|---|---|---|
| Land-use projections | | | |
| Population growth | Relatively low<br>(8.5 billion in 2050) | Low to high<br>(10 billion in 2050) | Relatively low<br>(8.5 billion in 2050) |
| Economic growth | High to medium<br>(284,565 GDP/PPP billion US$2005/yr in 2050) | Slow<br>(177,284 GDP/PPP billion US$2005/yr in 2050) | High<br>(360,926 GDP/PPP billion US$2005/yr in 2050) |
| Urbanization | High<br>(92% in 2050) | Low<br>(60% in 2050) | High<br>(92% in 2050) |
| Equity and social cohesion | High | Low | High |
| International trade and globalization | Moderate | Strongly constrained | High |
| Policy focus | Sustainable development | Security | Development, free market, human capital |
| Institution effectiveness | Effective | Weak | Increasingly effective |
| Technology development | Rapid | Slow | Rapid |
| Land-use regulation | Strong | Limited | Medium |
| Agricultural productivity | High | Low | High |
| Consumption & diet | Low growth, low-meat | Resource-intensive | Material-intensive, meat-rich diet |
| Mitigation policies in land use | Full | Absent | Absent |
| Bioenergy | High | Low | Lowest |
| Climate projections | | | |
| Carbon intensity | Low | High | High |
| Energy intensity | Low | Intermediate | High |
| Radiative forcing | Peak at 3W/m$^2$ before 2100 and declines | Stabilizes to 6W/m$^2$ in 2100 | Rising to 8.5 W/m$^2$ in 2100 |
| Concentration (p.p.m) | Peak at 490 $CO_2$ equiv. before 2100 then declines | 850 $CO_2$ equiv. (at stabilization after 2100) | >1,370 $CO_2$ equiv. in 2100 |
| Methane emissions | Reduced | Stable | Rapid increase |

**Table S2. Model description, metrics, and scenarios**

| Model | Description | Taxonomic scope | Metrics | Scenarios |
|---|---|---|---|---|
| AIM-biodiversity (Asia-Pacific Integrated Model – biodiversity) | A species distribution model that estimates biodiversity loss based projected shift of species range under the conditions of land-use and climate change. Species range shifts were projected under two commonly used dispersal assumptions: 'no' migration, which did not allow for species colonization and 'full' migration, which allowed for species colonization. Only the "no-migration" estimates were used. | Amphibians, birds, mammals, plants, reptiles | $S\alpha$<br>$S\gamma$<br>$H\gamma$ | Historical<br>Land use<br>Land use and climate |
| InSiGHTS | A high-resolution, cell-wise, species-specific hierarchical species distribution model that estimate the extent of suitable habitat (ESH) for mammals accounting for land and climate suitability. The model did not consider species colonization in this exercise. | Mammals | $S\alpha$<br>$S\gamma$<br>$H\gamma$ | Historical<br>Land use<br>Land use and climate |
| MOL (Map of Life) | An expert map based species distribution model that projects potential losses in species occurrences and geographic range sizes given changes in suitable conditions of climate and land cover change. The model considered range loss within the currently known distribution, and not the species colonization in this exercise. | Amphibians, birds, mammals | $S\alpha$<br>$S\gamma$<br>$H\gamma$ | Land use and climate |
| cSAR (Countryside Species Area Relationship) - iDiv | A countryside species-area relationship model that estimates the number of species persisting in a human-modified landscape, accounting for the habitat preferences of different species groups. | Birds | $S\alpha$<br>$S\gamma$ | Historical<br>Land use |
| cSAR-IIASA-ETH | A countryside species area relationship model that estimates the impact of time series of spatially explicit land-use and land-cover changes on community-level measures of terrestrial biodiversity. | Amphibians, birds, mammals, plants, reptiles | $S\alpha$<br>$S\gamma$ | Historical<br>Land use |
| *BILBI* (Biogeographic modelling Infrastructure for Large-scale Biodiversity Indicators) | A modelling framework that couples application of the species-area relationship with correlative generalized dissimilarity modeling (GDM)-based modelling of continuous patterns of spatial and temporal turnover in the species composition of communities (applied in this study to vascular plant species globally). | Vascular plants | $S\gamma$ | Historical<br>Land use<br>Land use and climate |

| Model | Description | Taxonomic scope | Metrics | Scenarios |
|---|---|---|---|---|
| PREDICTS (Projecting Responses of Ecological Diversity In Changing Terrestrial Systems) | The hierarchical mixed-effects model that estimates how four measures of site-level terrestrial biodiversity – overall abundance, within-sample species richness, abundance-based compositional similarity and richness-based compositional similarity – respond to land use and related pressures. | All | $S\alpha$<br>$I\alpha$ | Historical<br>Land use |
| GLOBIO | A modelling framework that quantifies the impacts of multiple anthropogenic pressures on biodiversity intactness, quantified as the mean species abundance (MSA) metric. | All | $I\alpha$ | Historical<br>Land use<br>Land use and climate |
| LPJ-GUESS (Lund-Potsdam-Jena General Ecosystem Simulator) | A big leaf model that simulates the coupled dynamics of biogeography, biogeochemistry and hydrology under varying climate, atmospheric $CO_2$ concentrations, and land-use land cover change practices to represent demography of grasses and trees in a scale from individuals to landscapes. | Not applicable | Bioenergy production<br>Food and feed production<br>Ecosystem carbon<br>Nitrogen retention | Historical<br>Land use<br>Land use and climate |
| LPJ (Lund-Potsdam-Jena) | A big leaf model that simulates the coupled dynamics of biogeography, biogeochemistry and hydrology under varying climate, atmospheric $CO_2$ concentrations, and land-use land cover change practices to represent demography of grasses and trees in a scale from individuals to landscapes. | Not applicable | Ecosystem carbon | Historical<br>Land use<br>Land use and climate |
| CABLE-POP (Community Atmosphere Biosphere Land Exchange) | A "demography enabled" global terrestrial biosphere model that computes vegetation and soil state and function dynamically in space and time in response to climate change, land-use change, $CO_2$ concentrations and N-input. | Not applicable | Ecosystem carbon<br>Timber production | Historical<br>Land use<br>Land use and climate |
| GLOBIO-E S | The model simulates the influence of various anthropogenic drivers on ecosystem functions and services. | Not applicable | Crop pest control<br>Nitrogen retention | Land use and climate |
| InVEST (Integrated Valuation of Ecosystem Services and Tradeoffs) | A suite of geographic information system (GIS) based spatially-explicit models used to map and value the ecosystem goods and services in biophysical or economic terms. | Not applicable | Coastal resilience<br>Pollination<br>Nitrogen retention | Historical<br>Land use and climate |

**Table S3. Description of land use categories in LUH2** (based on (*39*, *42*, *80*))

| | |
|---|---|
| forested primary land (primf) | natural vegetation that has never been impacted by human activities (agriculture or wood harvesting) and that is potentially forest; there is no transition to primary land from any other land cover categories |
| non-forested primary land (primn) | natural vegetation that has never been impacted by human activities (agriculture or wood harvesting) and is non-forest based on the LUH2 potential forest land layer; there is no transition to primary land from any other land cover categories |
| potentially forested secondary land (secdf) | natural vegetation that is recovering from previous human disturbance (either wood harvesting or agricultural abandonment) and is potentially forest; secondary land can never return to primary land |
| potentially non-forested secondary land (secdn) | natural vegetation that is recovering from previous human disturbance (either wood harvesting or agricultural abandonment) and is potentially non-forest; secondary land can never return to primary land |
| managed pasture (pastr) | land where livestock is known to be grazed regularly or permanently with some level of management activities, with low aridity and high population density |
| rangeland (range) | land where livestock is known to be grazed regularly or permanently, with high aridity and low population density; not managed except by grazing (i.e., no external inputs of pesticides or fertilizers, or fire/mowing) |
| urban land (urban) | areas with human habitation and/or buildings where primary vegetation has been removed |
| C3 annual crops (c3ann) | land where native vegetation has been removed and replaced with C3 annual crops; includes biofuel crops |
| C3 perennial crops (c3per) | land where native vegetation has been removed and replaced with C3 perennial crops; includes biofuel crops |
| C4 annual crops (c4ann) | land where native vegetation has been removed and replaced with C4 annual crops; includes biofuel crops |
| C4 perennial crops (c4per) | land where native vegetation has been removed and replaced with C4 perennial crops; includes biofuel crops |
| C3 nitrogen-fixing crops (c3nfx) | land where native vegetation has been removed and replaced with C3 nitrogen fixing crops; includes biofuel crops |
